# ‘Crafting Fishy News’: Framing and Attitudinal Positioning in English Newspaper Articles on Mahseer from Their Endemic Range

**DOI:** 10.1101/2025.05.22.655491

**Authors:** Prantik Das, V. V. Binoy

## Abstract

With the capacity to influence readers‘ knowledge, emotions, attitudes and behaviours, newspapers play a pivotal role in conservation. We studied framing strategies and attitude positioning in articles published from January 2000 to April 2024 on mahseer, a group of freshwater fishes (56 species), by English-language newspapers from 17 nations that fall within their distribution range. However, we had to restrict our analysis to 6 nations (Bangladesh, Bhutan, Nepal, India, Pakistan and Malaysia) and 6 mahseer species (*Tor khudree, T. putitora*, *T. remadevii*, *T. tor*, *N. hexagonolepis,* and *T. tambroides*) due to the limited availability of substantial number of articles from other focal nations. A major share (79%) of the articles came from India, and *T. putitora* was the species that received the maximum attention from the newspapers (39%). Our result revealed a clear demarcation between the South-East Asian nation of Malaysia and the countries from South Asia, as well as between articles on mahseers native to these two different geographical regions, in terms of headline and text framing, choice of messengers and images, and attitude positioning in the news items. Dailies from South Asia largely employed conservation, environmental challenges and re-stocking efforts frames and promoted nature-centric, ecological and religio-cultural attitudes. By contrast, newspapers from Malaysia and the articles covering mahseer from this nation, *T. tambroides* reflected utility-centric and recreational framing and attitudes emphasising economic aspects. In general, episodic framing was more common than thematic, and instances of incorporating visual representations (with India as an exception) and positioning information on mahseer in the persuasive sections (exception - Nepal) were limited across focal nations and species. Our study also revealed that GenAI based Large Language Model (LLM ChatGPT 4) and Natural Language Processing (NLP) based tools (Hugging Face transformers) can be utilised as effective complementary or collaborative research tools alongside traditional manual qualitative data analysis to comprehend the framing and attitude positioning in the conservation news. Our results may help contextualise the communication strategies for conserving mahseers in South and South-East Asia, and also elucidate dimensions that need to be made salient for strengthening the connection between media and society for this purpose.

## Introduction

Reliable information from trustworthy sources could influence stakeholders‘ perceptions and opinions on issues related to biodiversity and conservation (Walker et al. 2019; López-Baucells et al. 2023). Media along with disseminating news and knowledge regarding non-human species, ecosystems and the need for protecting them (Sen and Goswami 2022; Gessa et al. 2024; Trainotti et al. 2024), also play the role of an active conservation actor by ‗setting agenda‘ (Iyengar and Simon 1993; McCombs 2005) and guiding the thinking and actions of the public. For instance, projection of the values of a wild species by the media could promote its conservation (Abroms and Maibach 2008; Rust 2015; Walker et al. 2019; Sen and Goswami 2022; Trainotti et al. 2024; Gessa et al. 2024), while biased, inaccurate, exaggerated, or unscientific information may hinder the efforts to protect them and fuel human-wildlife conflicts (Jacobson et al. 2012; Muter et al. 2013; Chandelier et al. 2018; López-Baucells et al. 2023). Conservation news is not just a description of a species, ecosystem, events or policy. The headlines, tone of the text, visual or graphical contents (individual frames; Entman 1993) etc. are carefully chosen to highlight or soften certain aspects of the news by the authors and publishers keeping their own goals (increasing readership, policies and politics they support etc.; Papworth et al. 2015; Walker et al. 2019; Correa-Chica et al. 2024) and the interests of the spectators in mind. Hence, understanding how media outlines the conservation news, and the short and long term influence such information bring on the mind-set and behaviours of the audience heterogeneous in their knowledge and attitude is of great importance in developing effective conservation programmes and policies especially in the case of species or ecosystems spanning multiple political and cultural boundaries.

Different kinds of frames are used by the media to chaperon the audience not only in what to think about but also in how to deliberate. Although there is no universally accepted definition available for frames due to the complexity of the concept, ―selecting and highlighting some facets of events or issues, and making connections among them so as to promote a particular interpretation, evaluation, and/or solution‖ (Entman 2003) is the one popular amongst the scholars. The meaning generated by a frame is a product of four major component – the communicator, texts and images used, receiver, and culture, and along with the first two, action and the context are used as the framing tools (Ali and Hassan 2022) to make a news ‗noticeable, meaningful or memorable‘ (Entman 1993). Conventionally media uses either social or naturalistic frames while broadcasting information related to the environment or conservation. Social frames emphasise the human factors – behaviour, attitude, society, cultural norms etc. while natural frames describe the events more as a natural phenomenon. The conservation news could also be outlined as specific instances or individual events (episodic frames), or as an attempt to position an issue within the broader socio-cultural, ecological or environmental contexts (thematic frame; Iyengar 1991; Siemer et al. 2007). The articles following the latter framing strategy may include background information and probable solutions for the environmental challenges. Many authors consider the word, tones, writing style, linguistic and visual content chosen by the communicator to facilitate discouraging or encouraging information seeking and engagement by leveraging the emotional appeal also as frames (Nabi 2003; De los Santos and Nabi 2019).

Newspapers frame news to shape cognitive, affective and behavioural components of the attitude of receivers (known as attitude positioning; White 2012; 2025; Rodríguez and Hernández 2012; Ahmad and Fatima 2024) by using attention grabbing headlines, incorporating persuasive phrases and lexicons, quotes from the influential messengers etc. into the text and displaying images with the potential to elicit emotional responses. Along with summarising the essence of the article, headlines also help readers to choose items for detailed reading from the surplus of stories present on each page (Geer and Kahn 1993; Ifantidou 2023; Ecker et al. 2014). Meanwhile, textual devices such as metaphors, analogies, rhetorical questions, thought-provoking statements make news more compelling and influential. Adding ‗messengers‘ or ‗quoting sources‘ into the content also enhances the credibility of the news and promotes attitude positioning (Muter et al. 2013; Walker et al. 2019; Belanger 2022). Already existing trust towards the messengers such as politicians, scientists, government officials, etc. in the society is often seen translating into the significance of the news (Belanger 2022; Hollihan and Baaske 2022; Whiting et al. 2019). Furthermore, persuasive news including columns, editorials, opinions, commentaries, letters, features sections of the newspapers etc. written by the invited stakeholders, senior editors or experts from the field could also function as the attitude positioner (Benoit-Barné et al. 2021; Gessa et al. 2024; White 2025). With the advancement of the printing technology and popularisation of the modern tools of digital photography images that could attract eyeballs and encourage public support for conservation activities (Coleman 2010; Rodriguez and Dimitrova 2011; Uggla 2018; Shaw et al. 2022; Ison et al. 2024) have become an increasingly significant component of the conservation communication (O‘Neill 2013; 2020). Compared to the text being indexical, images can simplify information, reduce cognitive load and elicit strong emotional reactions (Joffe 2008; McInerny et al. 2014; O‘Neill 2020) and thus kindle faster public responses.

Although internet-based channels and social media are evolving as the popular sources of news and knowledge, newspapers, both in their printed and online versions, still maintain their reputation of ‗social influencer‘ in many nations. Middle-aged individuals and rural audiences continue to uphold newspapers as a relevant source of information (Martin and Copeland 2003; Bakker 2011; Martens et al. 2018; Agarwal and Sarkar 2022), and many considers this outlet as authentic and trustworthy in this era marked by the widespread mis-, dis- and mal-information (Martens et al. 2018; Agarwal and Sarkar 2022). Media framing and attitude positioning of the wildlife and conservation news has been extensively studied keeping print media (newspapers, magazines, journals, etc.) and their online versions as focal points all over the world (Jacobson et al. 2012; Muter et al. 2013; Shiffman et al. 2021). Unfortunately, much of the focus of such studies are terrestrial and marine species (Chandelier et al. 2018; Walker et al. 2019; Belanger 2022; Correa-Chica et al. 2024; Gessa et al. 2024; Ison et al. 2024) and the approach newspapers take towards the freshwater species remains largely unexplored. Although may be due to the low species detectability, restricted accessibility and research requiring huge amounts of resources, freshwater life-forms fail to attract the much-needed attention of the conventional media (Carrizo et al. 2017). However, studies focusing on the coverage of these species in conventional and social media could provide vital insights not on their availability, distribution, perceptions and attitudes of public and policy makers (Das and Binoy 2024) but also reflect various challenges faced by their habitat, an over exploited natural resource (Kolandai-Matchett and Armoudian 2020).

Mahseers, a group of iconic freshwater megafishes (out of the 56 known species many can grow above 30 kg) are increasingly becoming focal points of media and academic attention in the recent past due to their ecological, cultural and economic significances, multiple anthropogenic pressures experienced by their natural populations and restocking programmes implemented by various governmental and non-governmental agencies (Fig 1; Pinder et al. 2019; Sarma et al. 2022). Presence of these species is essential to ensure the health of riverine ecosystems and they hold significant values as food, source of livelihood and popular game fish. Mahseers are natives to 17 nations spanning across South, East and Southeast Asia known for their ecological, socio-cultural and economic diversity as well as for the noticeable variations within and across countries in the values attributed to various biotic and abiotic natural resources (Nautiyal 2014; Muchlisin et al. 2022; Redhwan et al. 2022). Many mahseers inhabit trans-boundary rivers spanning multiple countries and regions (Pinder et al. 2019; Sarma et al. 2022) and making the scenario further complex different localities of a single geographically large nation (such as India) has been shown to diverge in their response to mahseers (Das and Binoy 2024). Hence, understanding public attitudes, social narratives, exploitation practices and the broader socio-environmental issues affecting their habitats at different geographical scales and social structures is vital to come out with effective plans to conserve mahseers. Newspapers being the mirrors of the society analysis of the articles published from different nations holding mahseer populations may give many vital insights for the development of integrated and public involved population management strategies and policies to protect existing mahseer populations. Despite its potential to aid conservation of the mahseers no such study has appeared in the literature till date.

**Fig 1.**
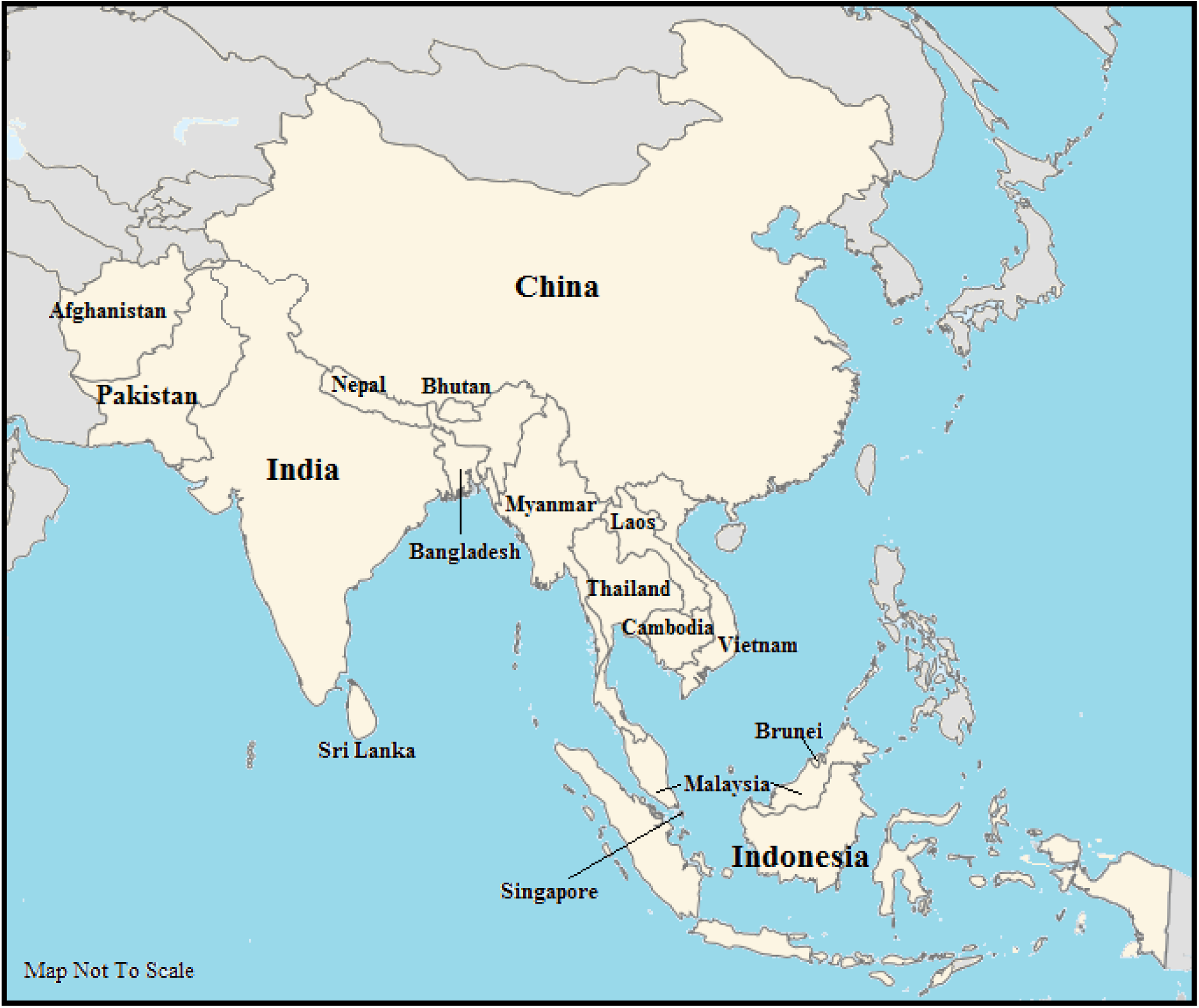
Endemic distribution range of the 56 known mahseer species across the globe. Map not to the scale.

Generative Artificial Intelligence (GenAI) based Natural Language Processing (NLP) tools such as Large Language Models (LLM), Multi-Natural Language Inference (MNLI)-based Transformers etc. have made processing and analysis of the enormous amount of text produced by the newspapers effortless and effective. Nowadays, these tools with the potential to improve speed, accuracy, efficiency and reliability of coding of the texts is being utilised as independent, complementary or collaborative research tools with traditional Manual Qualitative Data Analysis (MQDA) by many researchers (Johnson et al. 2023; Şen et al. 2023; Avinç and Yıldız 2024; Bijker et al. 2024; Christou 2024; Laajini and Tadjousti 2024; Perkins and Roe 2024; Prescott et al. 2024; Yan et al. 2024). However, this trend has not yet received the popularity in the field of conservation communication research focusing on aquatic species. Hence, we explored the following questions on framing and attitude positioning of the articles on different mahseers species published by the selected English language newspapers from 17 countries falling on their distribution range:

1. How did English newspapers in the focal nations framed headlines, textual contents and accompanying images, and engaged messengers/quoting sources in the articles on different species of mahseers?
2. How did these newspapers execute attitude positioning while reporting information on diverse mahseer species?
3. Could AI based tools ChatGPT 4.0 and Hugging Face transformers generate attitude positioning scores comparable to MQDA?

## Materials and Methods

### Data Collection

We generated a list of English newspapers available on the websites of the concerned Government Ministries of 17 countries spanning across South, East and South-East Asia from which the natural populations of mahseer are reported (Table 1). Audit Bureau of Circulation (ABC) and the BBC Media Guide were the source when this information was absent from the websites of the ministries (Brunei, Cambodia, China, Indonesia, Laos, Singapore, Thailand and Vietnam). From this list English dailies publishing both print and online editions were searched manually using a set of keywords related to mahseer (Supplementary Material; SM 1). Anglicized vernacular names of the focal species were not included as the keywords to avoid the potent ambiguity from regional variations. One week/fortnight free-trial available was utilised for collecting data for the newspapers demanding subscription for accessing the full article. From this database any country or species with less than ten articles published during the period January 2000 to April 2024 were excluded from further analysis (SM 2). The articles that contained the term mahseer but were not related to the fish (3 in number, e.g. Wild Mahseer Resort) were also discarded. The year of publication, headlines, category of news (editorial, columns etc), place of publication (including the name of states in the case of Indian newspapers), and the full text and images were extracted manually from each of the news articles shortlisted.

**Table 1.** Number of English newspapers analysed and the articles on mahseer collected from the 17 focal nations. The sources of these newspapers were the websites of the concerned Government Ministries (A), Audit Bureau of Circulation (B) and BBC Media Guide (C). All English newspapers available in these lists were searched for the news on mahseer keeping 2000 to 2024 (April) as the time period.

| Country | Number of English Newspaper Searched | Number of News Articles on Mahseer Obtained | Source of Newspapers |
| --- | --- | --- | --- |
| Afghanistan | 3 | 0 | A (Ministry of Information and Culture), C |
| Bangladesh | 6 | 12 | A (Ministry of Information and Broadcasting) |
| Bhutan | 4 | 20 | A (Ministry of Information and Communication), C |
| Brunei | 1 | 0 | C |
| Cambodia | 2 | 0 | C |
| China | 4 | 2 | B, C |
| India | 21 | 417 | A (Ministry of Information and Broadcasting), B |
| Indonesia | 3 | 3 | C |
| Laos | 2 | 0 | C |
| Malaysia | 3 | 37 | A (Ministry of Communications), B |
| Myanmar | 2 | 0 | A (Ministry of Information), C |
| Nepal | 3 | 10 | A (Ministry of Communication and Information Technology) |
| Pakistan | 6 | 31 | A (Ministry of Information and Broadcasting) |
| Singapore | 2 | 0 | B |
| Sri Lanka | 4 | 0 | A (Ministry of Mass Media) |
| Thailand | 3 | 4 | B, C |
| Vietnam | 2 | 0 | C |

### Analysis

The news collected were segregated based on the country of origin and mahseer species mentioned. The publications that did not specify species name were added to the group ‗mahseer‘-a sub-category under the species. Both category of news was subjected to the following analyses:

#### 1. Headline Framing

Manual coding (MQDA) of the headlines extracted from each news article was done using NVivo 14. The MQDA involved a five-step process - (i) familiarisation with data, (ii) coding, (iii) searching for frames (themes) by charting codes into frames, (iv) reviewing frames and (v) interpreting data with frames. To complement this analysis one of the popular LLM, ChatGPT 4.0 was used as the second coder. The headlines of the news articles categorised by the country of origin and species quoted, were individually uploaded to ChatGPT 4.0. The prompt — ―conduct a qualitative inductive thematic framing analysis on the attached raw data‖ — was used to generate framing data for each focal nation and mahseer species. These headlines were further subjected to sentiment analysis using the AFINN Lexicon (Nielsen 2011) and scores as positive, neutral and negative on an integer scale of -5 to +5 to were also generated (Hossain et al. 2021; Mori et al. 2025).

#### 2. News Content Framing

The text of each news article available in the two categories - countries and species - was subjected to the episodic versus thematic framing study (Iyengar 1991), followed by sentiment analysis, and inductive framing analysis for tracing out other framing strategies used. Here also the tools used for the analysis were MQDA (using NVivo 14), AFINN Lexicon and ChatGPT 4.0. The ‗persuasive‘ nature of each article was determined based on the segment/section of the newspaper in which it appeared (e.g. editorials, opinions, columns, commentaries, features and letters; Benoit-Barné et al. 2021; Gessa et al. 2024). Individuals or organisations quoted as the source of information (messengers/quoting source; Belanger 2022; Painter et al. 2024) or placed as the key actor in the programmes related to mahseer were also identified. These ‗messengers‘ were further segregated into the following categories - researchers, NGOs, government officials, journalists, celebrities (authors, musicians and photographers), recreational anglers, politicians, temple/shrine managers, resort manager/owners, military personnel, restaurant owners, fish farmers and local public. The category ‗government officials‘ included members of national, state and divisional biodiversity boards, officials of forest departments, fisheries/hatchery, pollution board, tourism departments and district administration.

#### 3. Visual Framing

Images present in the news articles coming under both nation and species categories were examined for their literal and denotative content using manual deductive thematic analysis (O‘Neill 2020). A total of five primary frames were selected following the literature available on the analysis of climate change and marine conservation related graphical representation on the print media (O‘Neill 2013; 2020; Ison et al. 2024). To this list - mahseer (animal), conservation action, conservation threat, landscape and portrait of stakeholder – we added two more frames: recreation and culinary considering the valuable food and game fish status of mahseer in different focal nations. The detailed description of the criteria used for segregating the images into the 7 categories mentioned is available in Table 2. A single coder manually examined each image and generated the visual framing data.

**Table 2.** Description of the images classified for each of the seven visual frames.

| Visual Frames | Description of Images |
| --- | --- |
| Mahseer (Animal) | Mahseer species are visible and the main subject in the picture. Images can be up-close of showing mahseer in its habitat. |
| Conservation Action | People engaging in conservation activities like research, monitoring, awareness/outreach, breeding and release, hatchery, court rulings and policies. These images place the activity at the centre of the photo and not the person. |
| Conservation Threat | Negative impact and threats on mahseer and their habitat. They can include threats like river pollution, hydel projects and hydro-power units on dams, climate change, river linking, sand/gravel/boulder excavation and mining, river bed mining, human impacts of killing and over-fishing, impact of invasive alien species (IAS). They place the threat at the centre of the image and not the person. |
| Landscape | Pictures showing wide landscape views of an area of land or water. Includes images of rivers, forests and river banks. |
| Portrait of Stakeholders | Images include stakeholders looking at and posing for the camera. Stakeholders include government officials, researchers, members of NGOs, anglers, politicians, natives. |
| Recreation | Images that focus on recreational angling of mahseers and ecotourism related to angling, travel and leisure. |
| Culinary | Images that focus on mahseers as a food fish. They may include pictures of specific mahseer dishes in restaurants. |

#### 4. Attitudinal Positioning

The potential of each news item to influence cognitive, affective and behavioural dimensions of the readers‘ attitude was estimated using 3 different methodologies. First of all a list of 9 attitudinal categories was developed referring to important ecological attitude studies (Kellert 1985; 1991; Schultz 2000; 2002; Salazar et al. 2022; Table 3). Since single news could impact multiple elements of the attitude, every news article collected was tested using MQDA, ChatGPT-4 and Multi-Natural Language Inference (MNLI) model for its contribution to each of the nine different attitudinal categories. Here also we sorted the articles in accordance with the mahseer species mentioned and nations of publication.

a. MQDA: Every focal articles were read by a coder and attitudinal scores were assigned to each of the 9 categories using ordinal numerical scales, 0, 0.5, and 1 (0: no attitude expressed. 0.5: attitude present but subtle and not dominant. 1: attitude dominant and strongly expressed; Green et al. 2015).
b. b). *ChatGPT:* Using a prompt ―give scores to the uploaded data using the nine defined attitude categories on a decimal scale of 0 to 1,‖ the content of the all articles were tested individually using ChatGPT-4. Here also the sum of the scores was generated as done in the case of MQDA.
c. c). *Zero-Shot Classification with Hugging Face Transformers:* Hugging Face transformers framework using the BART-large Multi Natural Language Inference (MNLI) model, a pre-trained zero-shot classifier pipeline (Hugging Face n.d.) was employed to detect the attitude positioning on a decimal scale of 0 to 1 of all the articles shortlisted. Here also the predefined nine categories were used as the reference. This model‘s flexibility to handle novel categories without additional training made it particularly suitable for our analysis (Imran 2024; Hugging Face n.d.). Hugging Face transformers were not unlisted in the framing studies mentioned in the above sections as they are primarily designed for quantitative tasks, such as scoring and classification relying on syntactical patterns and word frequencies (Hugging Face n.d.). Framing analysis requires a deeper understanding of contextual and background knowledge, such as the ecological and socio-cultural significance of mahseers in different nations, not explicitly stated in the text, which was not a forte of MNLI models.

**Table 3.** News articles on mahseer were explored for the 9 different types of environmental attitudes to know the attitude positioning.

| S. No. | Attitudes | Description |
| --- | --- | --- |
| 1. | Nature-centric | Affection, care and closeness with nature |
| 2. | Ecological | Concern for environmental degradation and habitat destruction, conservation issues and solutions |
| 3. | Empathetic | Affection and concern towards mahseers with a focus on their feelings, emotions and experience |
| 4. | Moralistic | Concern for the right and wrong towards mahseer and nature |
| 5. | Research-oriented | Research and scientific findings / studies |
| 6. | Religio-cultural | Religious, cultural or symbolic representation of mahseers |
| 7. | Utility-centric | Mahseers used as food and for material products |
| 8. | Recreational | Sporting and recreational activities |
| 9. | Negative | Feelings of hate and/or indifference towards mahseers |

For every focal nation and species, a single value for each of the 9 different attitude categories was generated by averaging the scores given to the individual articles {Table 4, 5; SM 3 (a-d); SM 4, 5}. These scores obtained from MQDA, ChatGPT-4 and Hugging Face transformers analysis were compared using Kruskal-Wallis rank sum test across countries and species. Data coding and organisation were performed manually using NVivo 14. The sentiment analysis, statistical tests and data visualisations were conducted on R version 3.6.1 and RStudio version 4.4.1. using ‗ttm‘, ‗syuzhet‘, ‗ggplot2‘, ‗reticulate‘, ‗reshape2‘ packages. KyPlot 6.0 (KyensLab Inc., Tokyo, Japan) was also used for the data visualisation.

**Table 4.** Proportion of the mahseer related news following episodic and thematic framings published from six countries (a) and centring on focal species (b). Methodologies used were MQDA and ChatGPT-4 analysis

| (a) Countries | Episodic (%) |  |
| --- | --- | --- |
|  | MQDA | ChatGPT |
| Bangladesh | 58.34 | 50 |
| Bhutan | 80 | 80 |
| India | 70.60 | 71.56 |
| Malaysia | 81.08 | 83.78 |
| Nepal | 70 | 70 |
| Pakistan | 74.20 | 70.96 |
| (b) Mahseer Species | MQDA | ChatGPT |
| *T. khudree* | 69.35 | 66.13 |
| *T. putitora* | 71.36 | 71.83 |
| *T. remadevii* | 62.50 | 62.50 |
| *T. tambroides* | 80.95 | 76.20 |
| *T. tor* | 63.15 | 68.42 |
| *N. hexagonolepis* | 82.45 | 80.70 |
| Mahseer | 70.90 | 69.40 |

## Results

A search of the 71 English newspapers listed from 17 focal nations produced only 540 news articles containing the term ‗mahseer‘ published in the last 24 years (from January 2000 to April 2024). No such news was available from Afghanistan, Brunei, Cambodia, Laos, Myanmar, Singapore, Sri Lanka and Vietnam, while China, Indonesia and Thailand published less than five pieces during this period (Supplementary Materials SM 2). India, the country that published the highest number of news articles on mahseer (417 articles) was far ahead of the next nations Malaysia (37 articles) and Pakistan (31; Table 1) in the list. Till 2008, each of the five nations shortlisted for the further study had an yearly publication of 0 to 4 articles on mahseer (Fig 2a, b). However, India witnessed a gradual increase in such news after this year, which spiked to a range of 30-50 articles per year past 2015. An analysis of the state wise publication of the news related to the focal fishes in India revealed Karnataka (27%), Nagaland (12.8%) and New Delhi (8%; Fig 3) as the main contributors. *T. putitora* was the mahseer species maximum represented in the newspapers across all nations (39%), while *T. tor* (3.48%) was the lowest to be covered (Fig 4).

**Fig 2.**
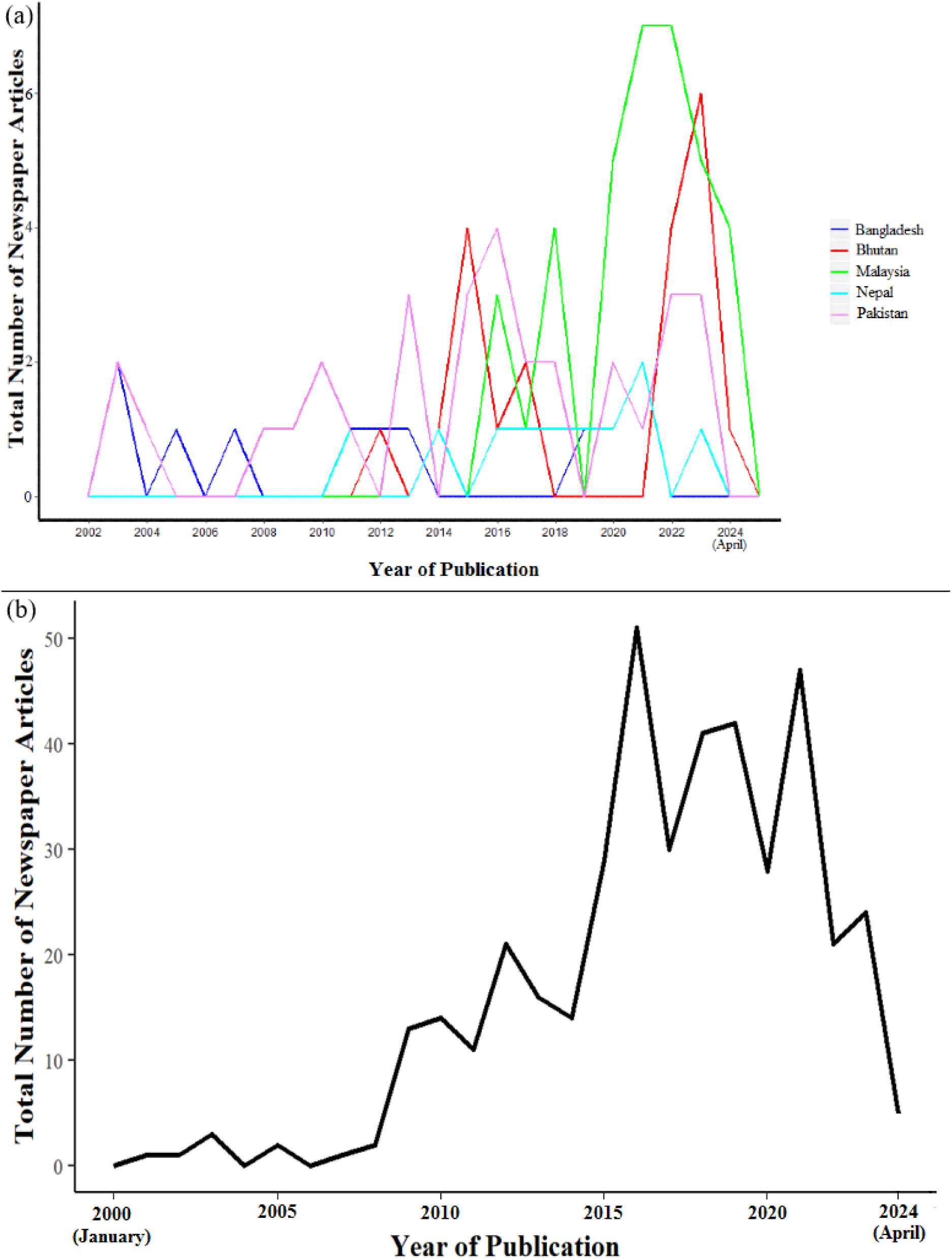
Articles published on mahseer by the English newspapers from five focal nations - (a) Bangladesh, Bhutan, Malaysia, Nepal and Pakistan and (b) India. The time period considered was from January 2000 to April 2024.

**Fig 3.**
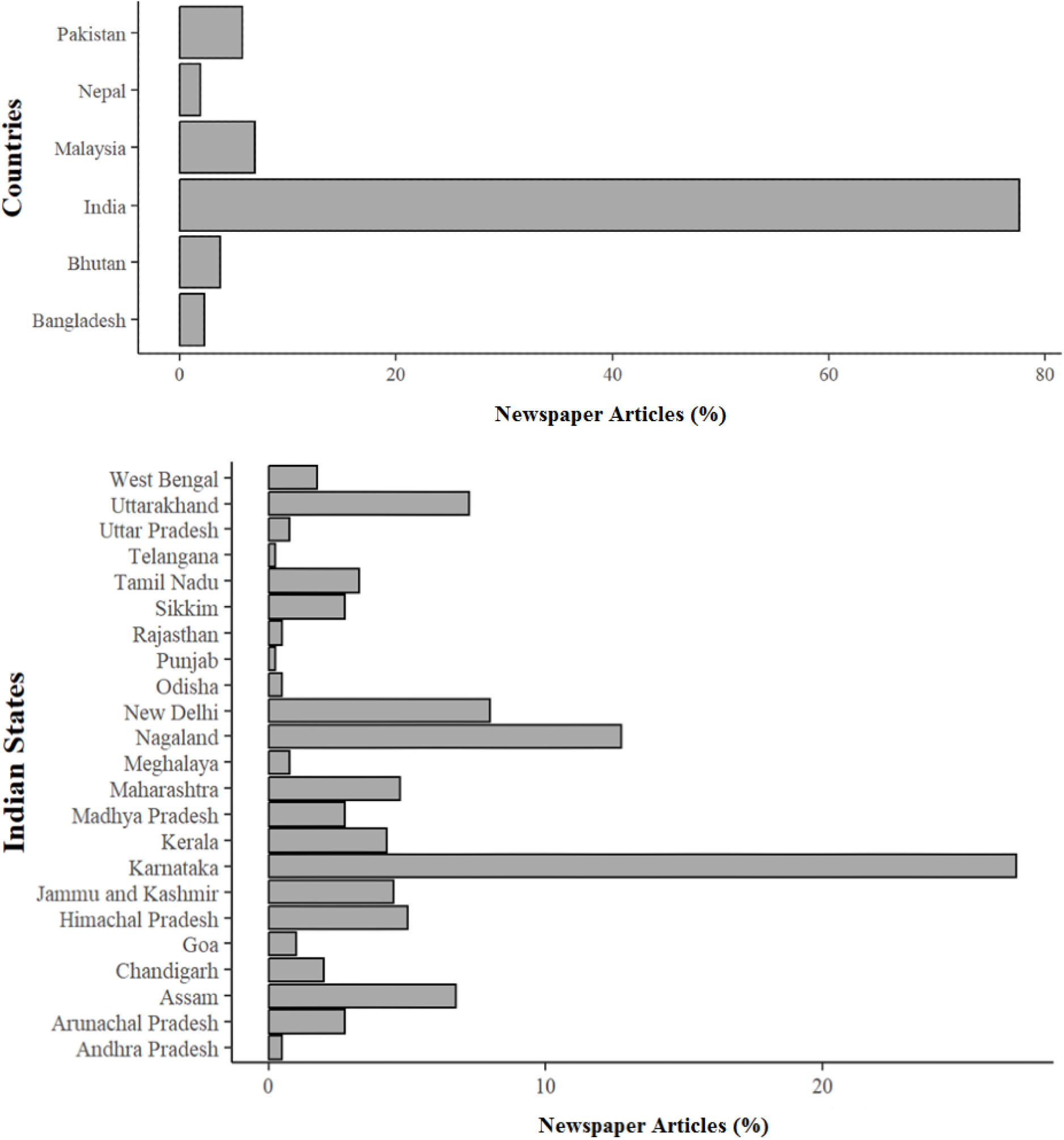
Articles on different mahseer species published by the English newspapers from (a) the six focal nations and (b) different states and union territories of India.

**Fig 4.**
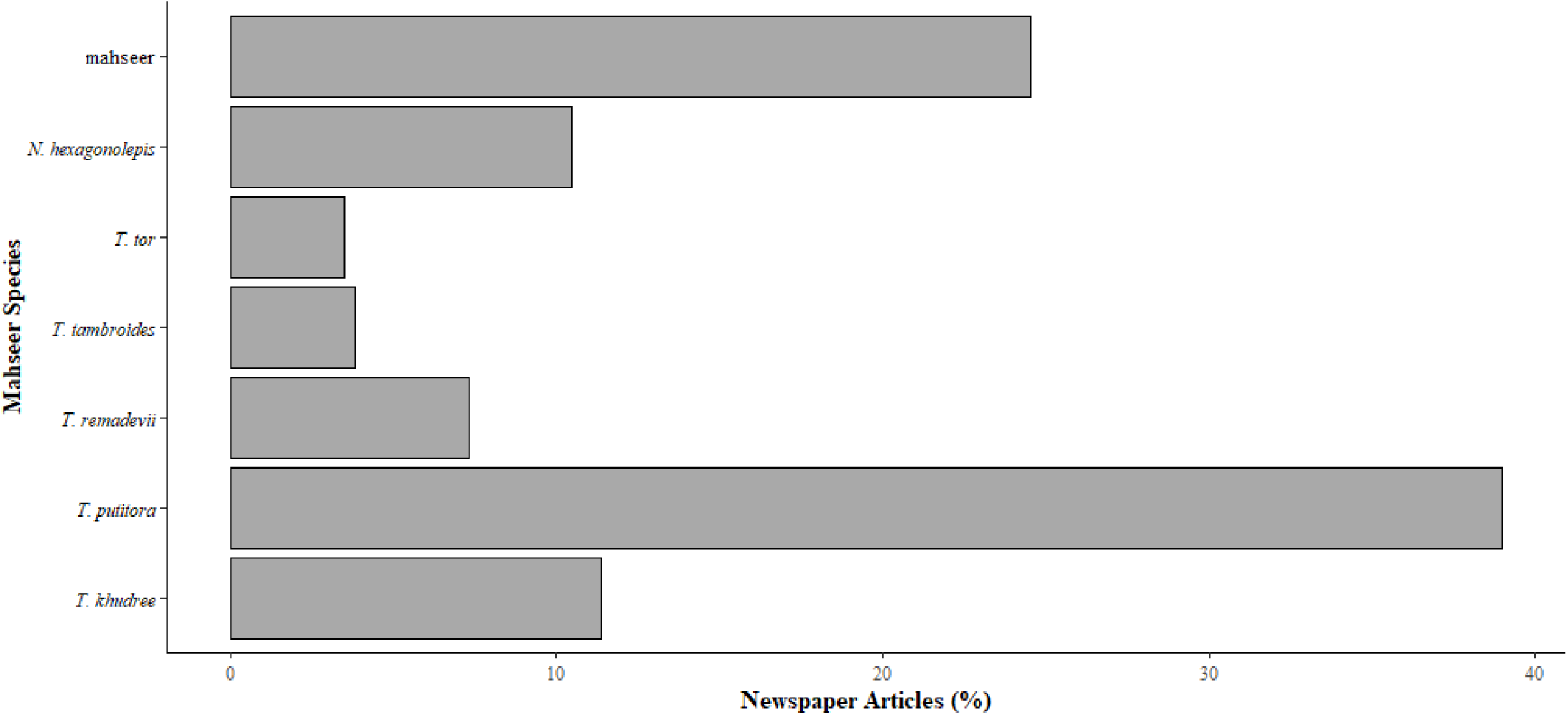
Articles on different mahseer species published by the English newspapers from the six focal nations. Only the species with more than 10 articles were considered for the study.

### 1. Headline Framing

MQDA revealed 13 frames - conservation, restocking, challenges faced by the mahseers, policy, law enforcement, education and awareness, science and research, recreational angling, cultural significance, human-human conflict, culinary, farming and revenue and travel and tourism - from the headlines of the news on mahseer published across the focal nations. The headline framing analysis using ChatGPT 4.0 resulted in 14 frames out of which 11 were consistent with MQDA. The ChatGPT divided MQDA frames ―challenges faced‖ into ―environmental threats‖ and ―habitat destruction effects‖ and clubbed ―policy‖ and ―law enforcement‖ to ―policy, law and governance‖. The focal nations diverged in framing the headlines of the news on mahseers. Headlines from Pakistan (P) and Bangladesh (B) were centered more on conservation of these megafishes (P, 25.61%; B, 30.24%), environmental challenges faced them (P, 25.61%; B 69.76%) and efforts done to restock their populations (23.26%). By contrast Nepali (36.31%) and Malaysian (32.11%) newspapers gave weightage to travel and tourism, with the former also highlighting the recreational angling (20.75%) mahseer farming and revenue from it (21.25%; Fig 5a). Another important observation was the lack of headlines containing frames on recreational angling in Bhutan, a country where this activity is popular, and science and research in Bangladesh. Although a major share of mahseer news analysed during the present study came from India, no frames were found dominating the headlines. However, though few in number, only news from this nation had headlines with a policy (5.82%) and human – human conflict (3.02%) angle (Fig 5a). Challenges faced was the most prominent frame of the headlines of the news focusing on mahseer species *T. khudree* (53.14%)*, T. remadevii* (66.67%)*, T. tor* (46.15%) and *T. putitora* (30.82%). Contrastingly, headlines of news articles on *T. tambroides* and *N. hexagonolepis* mostly featured topics related to their farming and revenue generation (40.02% and 28%; respectively Fig. 5b). Conservation, stocking, education and awareness, science and technology frames were totally absent in the case of *T. tambroides.* Though this species is a delicacy in Malaysia, culinary related headlines keeping them in focus were also not available. The sentiment analysis of the headlines from all six countries and focal mahseer species using the AFINN Lexicon resulted in a neural score of ‗0‘.

**Fig 5.**
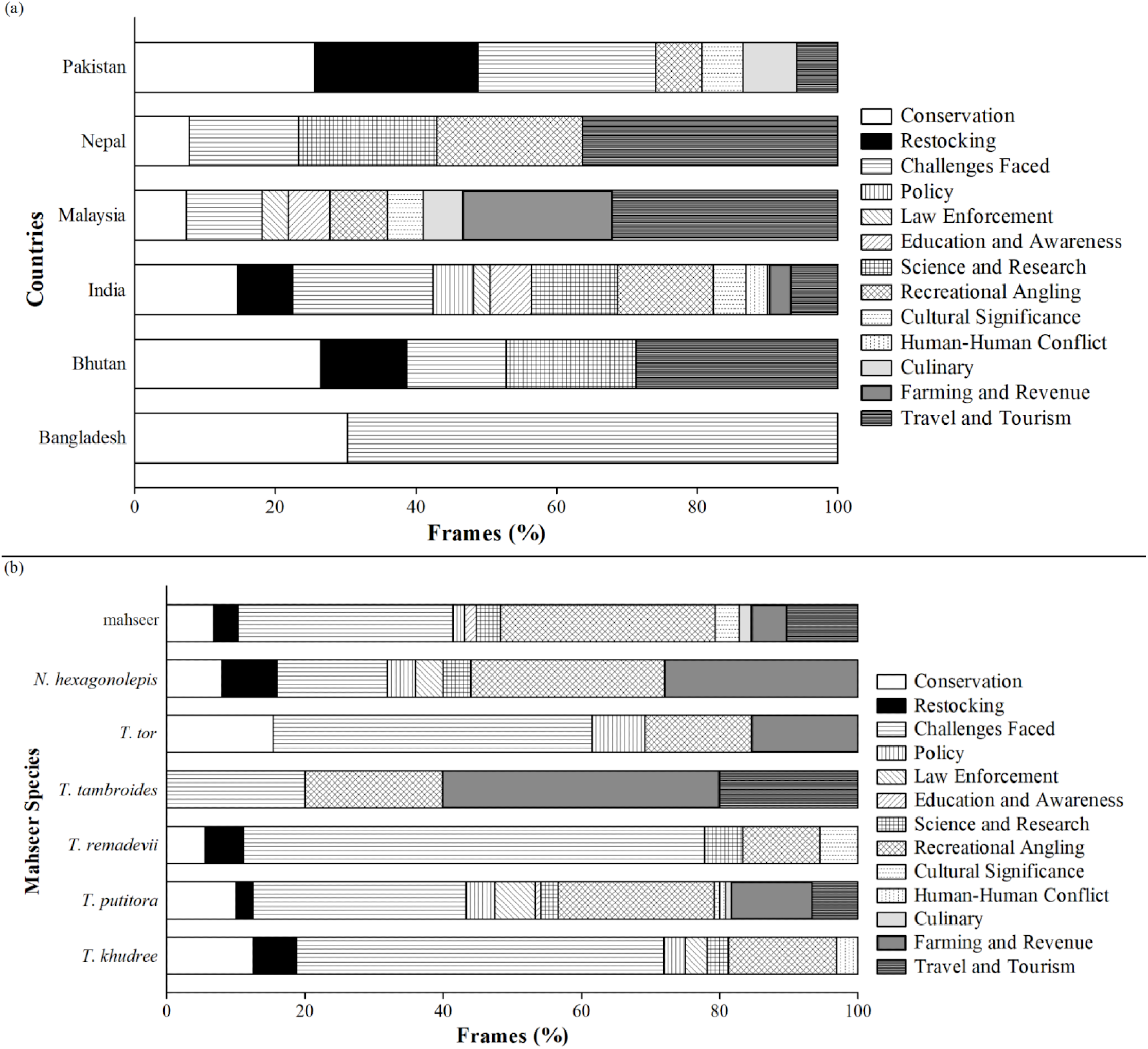
Distribution of different frames in the headlines of mahseer related news published (a) from six countries and (b) centring on focal species.

### 2. News Content Framing

Analysis of the content of the news articles produced 13 and 15 frames from MQDA and ChatGPT 4.0‘s thematic analysis respectively. This difference was the result of the latter segregating an MQDA frame ―challenges faced‖ into three: ―environmental threats,‖ ―habitat destruction impacts,‖ and ―extinction risks‖. In addition to the 12 frames that emerged out of the headlines, study of the news text produced one more, ―illegal fishing‖ (practices like dynamiting, poaching and destructive fishing). Meanwhile the ‗law enforcement‘ frame appearing in the headlines was absent in the contents. Hence a list of 13 frames was used for further analyses.

The nations from South Asia (Bangladesh, India, Nepal and Pakistan), framed the news mainly around challenges faced, restocking and conservation, while emphasis was given to the economic aspects (travel and tourism, culinary, farming and revenue) by Malaysia. Additionally, very little news on science and research (1.55%), as well as education and awareness (3.10%) on mahseers came out from this South-East Asian nation (Fig. 6a). In comparison to other countries, Bangladesh (13.51%), Nepal (11.36%) and Bhutan (11%) published news emphasising the cultural value of these large bodied freshwater fishes. Illegal fishing reported by Nepali (13.63%) and Pakistani (12.28%; Fig 6a) media and total absence of the information on restocking in the articles published by the former raises concerns for the mahseers in these countries. The focal countries‘ framing style reflected on the mahseer species they harbour. The frames related to conservation, challenges faced and recreational angling received considerable attention in the news stories covering *T. khudree, T. putitora* and *N. hexagonolepis* and *T. tor* the species reported from South Asia. News on *T. remadevii,* another mahseer from the same region also followed the same trend, but recreational angling news was less for this species. *T. tambroides,* the mahseer species found in Malaysia, was mentioned mainly in the articles centred on culinary and travel and tourism and gave least attention to challenges faced (6.02%), education and awareness (3.61%) and restocking (2.41%). Sadly, the science and research remained the minimum represented frame in articles on *T. tambroides* (1.20%; Fig. 6b). By contrast *N. hexagonolepis* received a relatively higher number of education and awareness (13.34%) frames.

**Fig 6.**
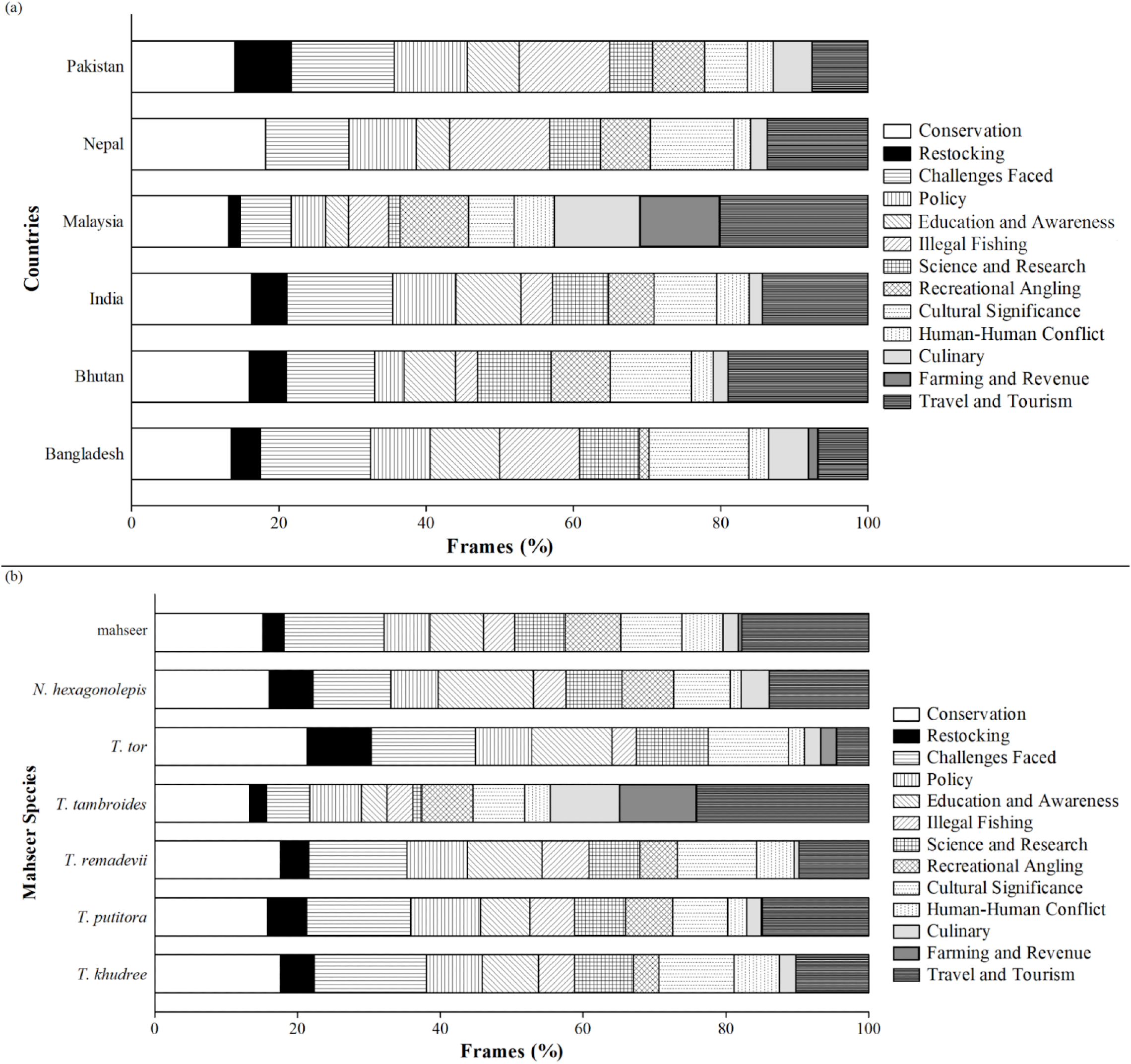
Distribution of different frames in the content of mahseer related news published (a) from six countries and (b) centring on focal species.

#### 2.1. Episodic vs. Thematic Content Framing

Mahseer news across the six countries found predominantly (more than 70%) employing episodic framing, except in Bangladesh (58.33%). Notably, the Malaysian (81.08%) and Bhutanese (80%; Fig 7a) dailies leaned heavily on this kind of framing. The repetition of the same analysis keeping various mahseer species in the focus also revealed a similar trend - around 60% of the articles available on each species were episodic in nature. Eighty percent of the news on mahseer from Malaysia (*T. tambroides,* 80.95%; Fig 7b) and species *N. hexagonolepis* (82.45%; from Bhutan, India and Nepal) showed episodic framing.

**Fig 7.**
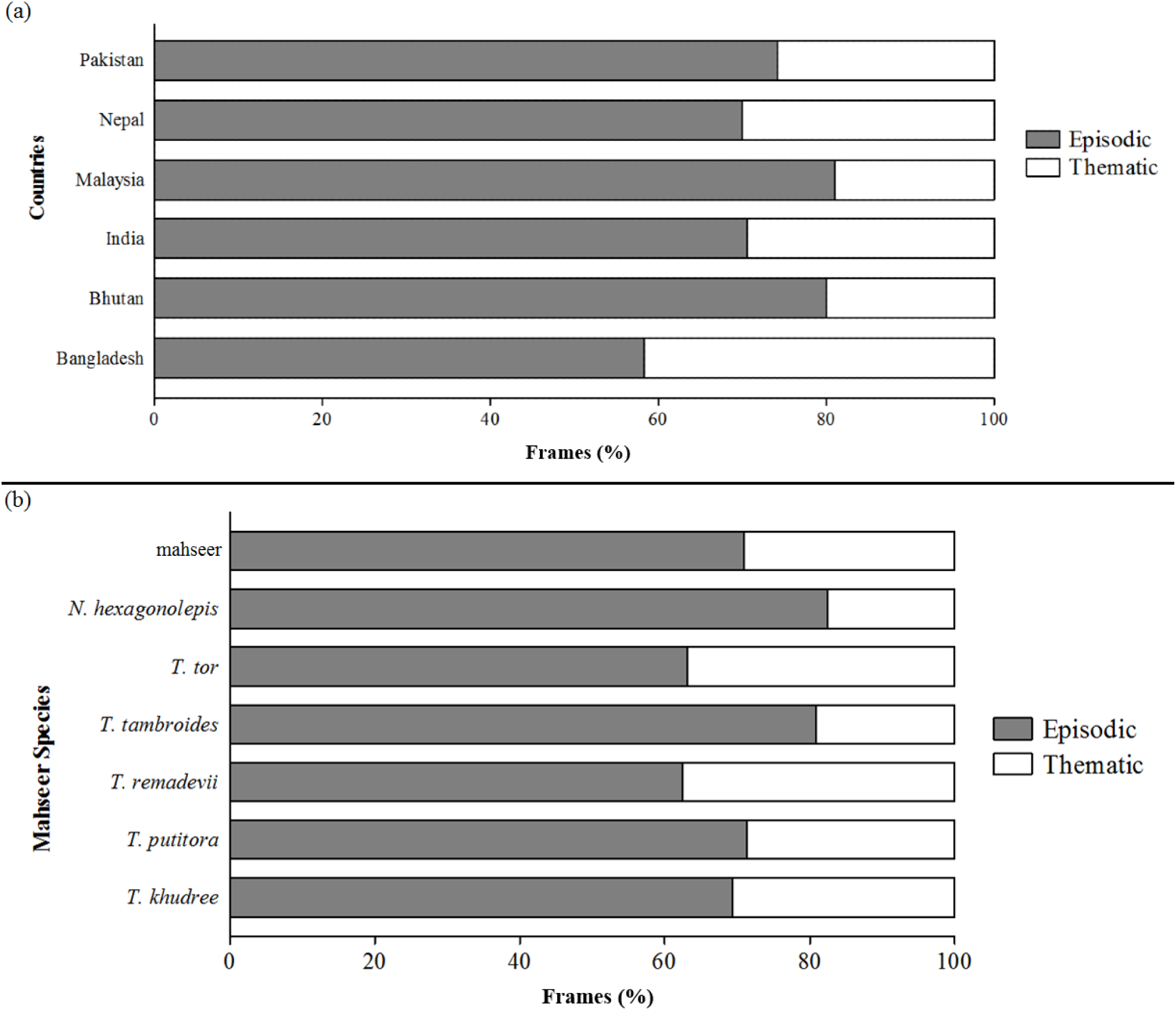
Usage of episodic and thematic framed in the content of mahseer related news published (a) from six countries and (b) centring on focal species.

#### 2.2. Sentiment Analysis

The sentiment analysis of the news content using AFINN Lexicon resulted in a neutral score of ‗0‘ for nations (Bangladesh, Bhutan, India, Nepal and Pakistan) and species (*T. khudree, T. putitora, T. remadevii, T. tor, N. hexagonolepis* and mahseer category). However, articles mentioning *T. tambroides* received a positive score of ‗+2‘ and the articles published from their home country Malaysia a ‗+2‘.

#### 2.3. Persuasive News

Two neighbouring nations from South Asia, Nepal and Bangladesh stood out in publishing persuasive sections such as editorials, columns, letters, opinion pieces, etc. on mahseer. Nearly 60% of the news published by the former fell in this section while the latter had 0 items in this category (Fig 8a). A good share of the news stories reporting *T. tambroides* (38.10%) had adopted a persuasive approach (Fig 8b).

**Fig 8.**
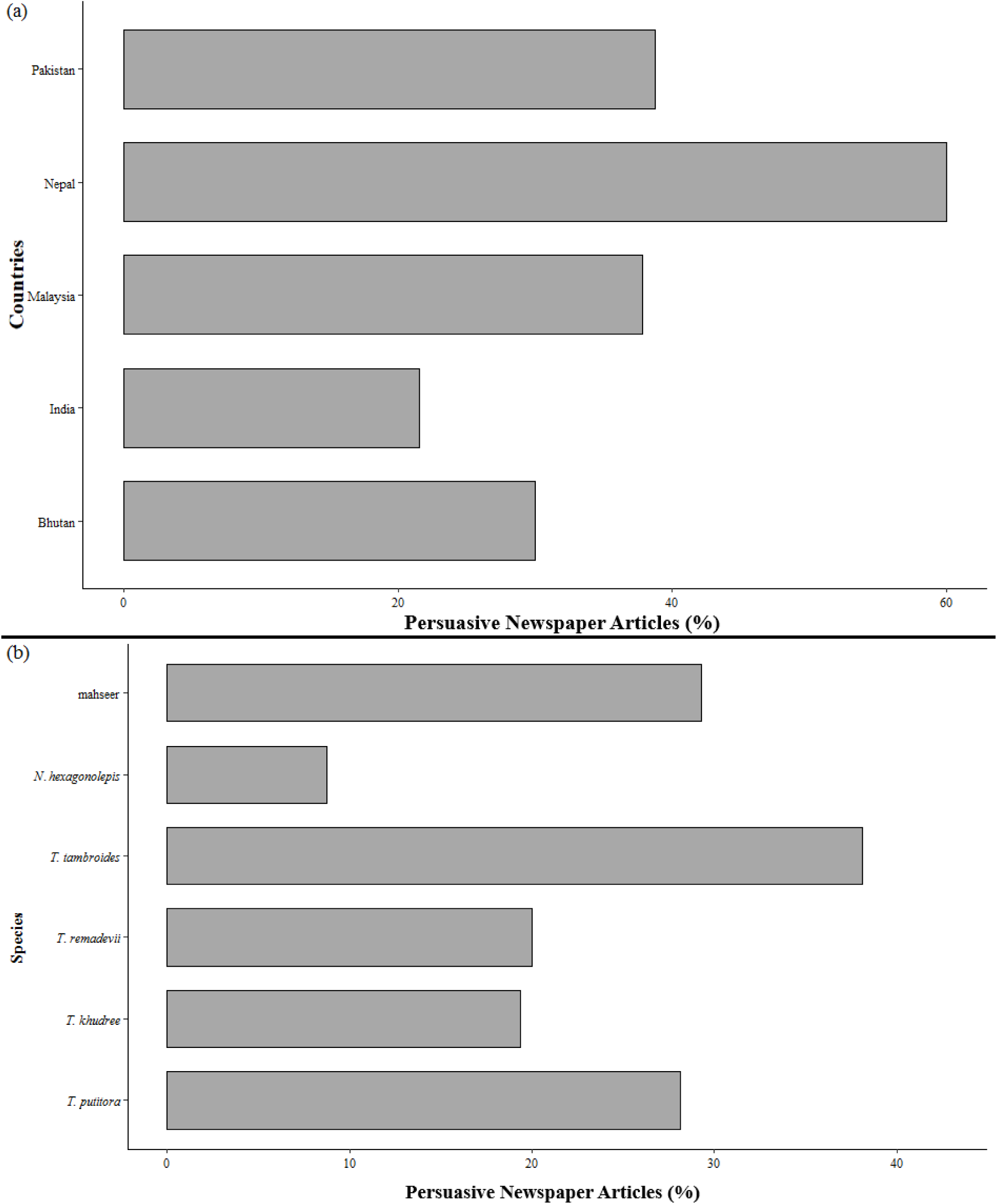
Persuasive news articles on mahseer published 9a) from six countries and (b) centring on focal species.

#### 2.4. Messengers/Quoting Sources

The individuals and organisations quoted in the news considerably varied, ranging from local people to politicians and celebrities amongst the countries studied. Interestingly, Bangladesh had the lesser diversity of the messengers in the mahseer news. Government officials were frequently cited in the news from Pakistan (60%), Bhutan (44.45%), Bangladesh (40%), India (35.42%) and Malaysia (34.61%) while recreational anglers decorated this position in Nepal (36.36%). In Bangladesh, researchers received exceptional attention from the media (40%). Although recreational angling of mahseer is popular in Bangladesh, Pakistan and Bhutan very few articles quoting anglers were published from these nations. Reflecting mahseer‘s religious significance, though small in number, temple and shrine managers were quoted exclusively in the articles published from Pakistan (2.50%) and India (0.58%). Fish farmers (15.38%; Fig 9a) and resort and restaurant owners (3.84% each) together constituted a noticeable group of messengers in Malaysian news articles. News from Nepal had a unique feature - celebrities and military personnel sharing their views on these important fishes. Media from all nations, except Nepal, dedicating a good amount of space to publish comments of the local people on mahseer, point towards a positive trend in their approach (Fig 9b).

**Fig 9.**
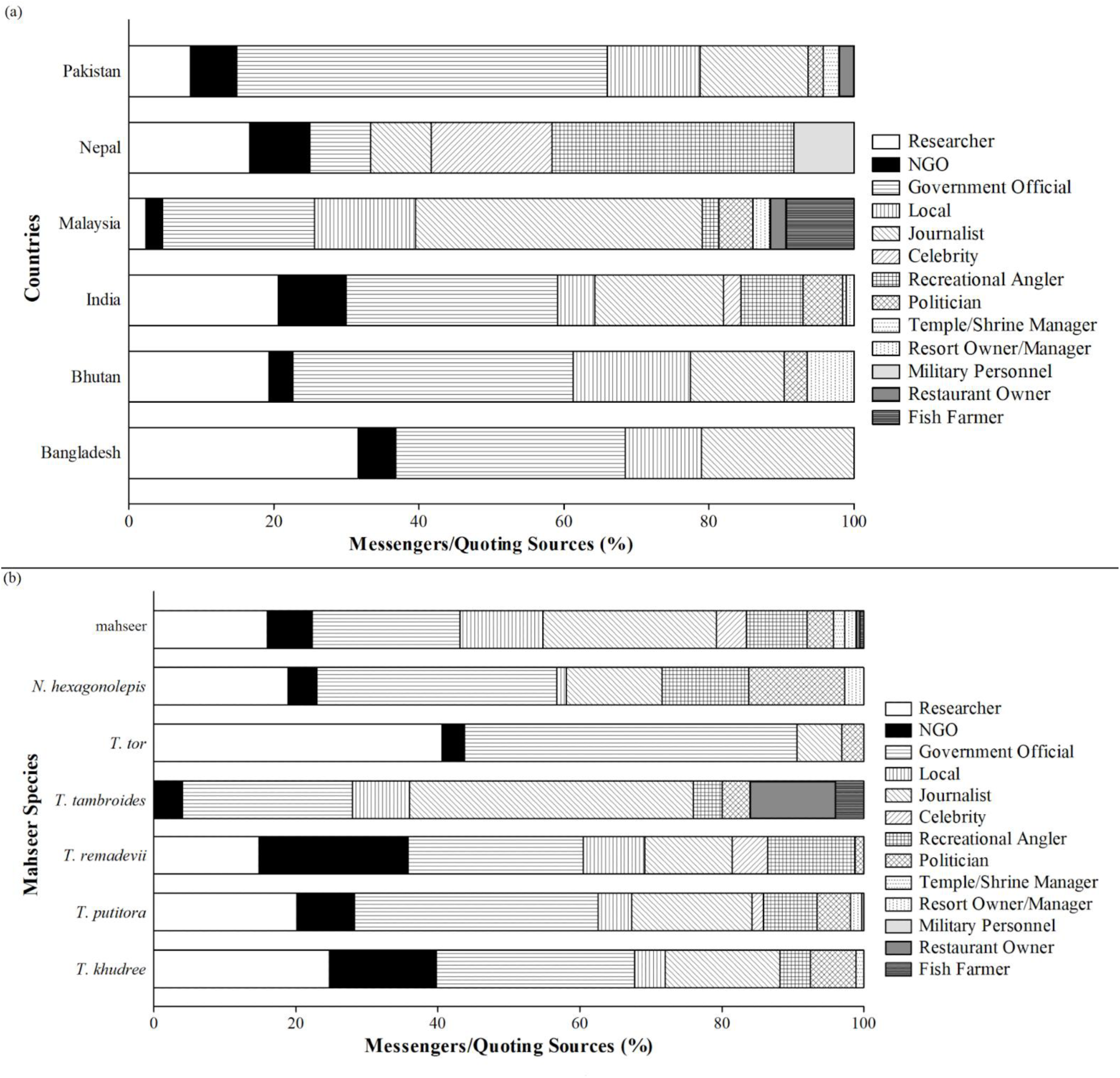
Messengers/Quoting sources present in the news articles on mahseer published (a) from six countries and (b) centring on focal species.

Government officials were prominently quoted in the articles on all mahseer species (*T. tor* 50%; *T. putitora* 41.28%; *T. khudree* 33.34%; *T. remadevii* 28.17%; *N. hexagonolepis* 39.06%, *T. tambroides* 40% and ‗mahseer‘ 27.46%). Least diversity in the stakeholders cited was for *T. tor,* and here the sources were dominated by researchers (43.34%). Although news on all species except *T. tor* (no articles) contained the perspectives of localites, ‗mahseer‘ category scored relatively high for this category (15.50%). Articles on *N. hexagonolepis* contained mentions of more politicians (15.62%). No researcher and very few NGOs (6.67%) were seen quoted in the case of *T. tambroides* (Fig 9). News on a recently described species *T. remadevii* (2018), the ‗mahseer‘ (the category where no specific species was mentioned) and *T. putitora* contained the word of celebrities. Meanwhile, fish farmers (20%) and restaurant owners (6.67%) were found only in the stories focusing on *T. tambroides*.

### 3. Visual Framing

India stood out as the country with highest number of visual representations (73.76%) in the mahseer news. Next country in this list, Malaysia had only 13.30% of the articles with images. News coverage from Pakistan (6.08%), Bhutan (3.80%) and Nepal (3.04%), and the articles describing *T. remadevii* (9.88%), *T. tor* (2.28%) and *T. tambroides* (6.46%) featured only a small number of images and were therefore excluded from further analyses. Noticeably, all articles collected from Bangladesh were devoid of any visual representation. Further exploration revealed no image category dominating others, neither in the case of India nor focal species. Landscapes were the most prevalent (31.37%) frames in the articles from Malaysia (Fig 10a). Across species, *T. putitora* received high visual coverage (32.32%), followed by *T. khudree* (13.30%), *N. hexagonolepis* (11.02%) and ‗mahseer‘ (22.81%). Conservation action (*N. hexagonolepis,* 30.77%), landscape (mahseer, 27.11%; *T. khudree,* 25%), recreation related (*N. hexagonolepis,* 19.23%; *T. putitora*, 20.48%) were the image types in noticeable quantity observed in the case of different mahseer species. The culinary frame appeared exclusively in the ‗mahseer‘ category articles (8.47%; Fig 10b).

**Fig 10.**
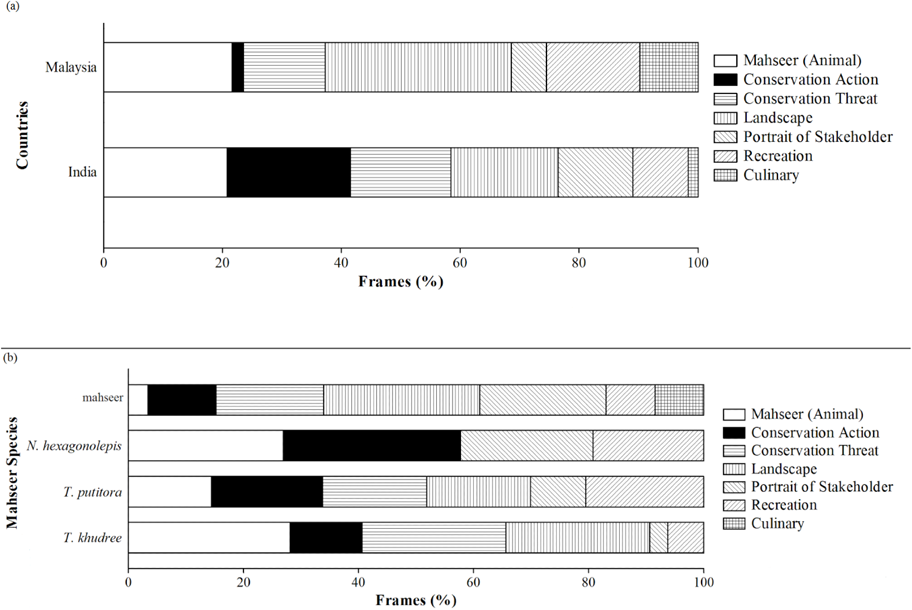
Visual frames of the images present in the news articles on mahseer published (a) from two countries and (b) centring on focal species. Only the countries and species whose news articles contained images in at least 10% of the total articles were considered for the study.

### 4. Attitudinal Positioning

The three different scores for each of the nine attitudinal categories determined using MQDA, ChatGPT and Hugging Face transformers (using zero-shot classifier), was not significantly different either for focal countries or species (Kruskal-Wallis rank sum test; Table 5; 6). Hence, inter and intra nation (as well as species) variation in the attitude positioning were explored using the scores obtained through MQDA (Kruskal-Wallis rank sum test followed by *post hoc* analysis using Dunn test with Bonferroni method). Except ‗empathetic (SM 5) and negative‘, all other attitudinal categories revealed significant inter-country differences (Table 7; Fig 11). Further analysis revealed a segregation of the nations from South Asia on one side and Malaysia on the other in the case of many attitudinal categories studied. For instance, Malaysia had significantly less number of mahseer news articles with ecological (in comparison to Bangladesh, India and Pakistan; SM 4), moralistic (India and Bangladesh; SM 6), research-oriented (Bangladesh, Bhutan, India and Pakistan; SM 7) and nature centric (Bhutan and India; SM 3) attitude positions. However, the recreational attitude was distinctly observable in Malaysian articles compared to Bangladesh, Bhutan, India and Pakistan (SM 10). India scored significantly higher in religio-cultural (than Bangladesh and Bhutan; SM 8) and nature-centric (Bangladesh and Pakistan; SM 3) attitudes, while Pakistan dominated moralistic (than Bhutan, and India; SM 6). Mahseers being exploited as a food source, the utility-centric attitude was significantly more prevalent in the news coverage from Bangladesh, Pakistan and Malaysia than in the other focal countries (SM 9). In comparison to Bangladesh, Nepal and India published significantly higher numbers of articles with nature-centric and recreational attitudes, respectively.

**Table 5.** Comparison of the three different scores for each of the nine attitudinal categories determined using MQDA, ChatGPT and Hugging Face transformers (using zero-shot classifier). Statistic used was Kruskal-Wallis rank sum test (df=2); * = *p* < 0.05, ** = *p* < 0.01, *** *p* < 0.005.

| **Attitudes** | **Bangladesh**<br>($\chi^2$) | **Bhutan**<br>($\chi^2$) | **India**<br>($\chi^2$) | **Malaysia**<br>($\chi^2$) | **Nepal**<br>($\chi^2$) | **Pakistan**<br>($\chi^2$) |
| --- | --- | --- | --- | --- | --- | --- |
| Nature-centric | 0.43 | 0.25 | 1.12 | 0.26 | 0.64 | 0.42 |
| Ecological | 0.31 | 0.18 | 0.65 | 0.64 | 0.12 | 0.14 |
| Empathetic | 1.04 | 1.10 | 0.98 | 1.56 | 0.76 | 1.09 |
| Moralistic | 0.83 | 0.75 | 0.82 | 0.24 | 0.35 | 0.51 |
| Research-oriented | 2.87 | 1.98 | 1.23 | 1.97 | 1.76 | 2.04 |
| Religio-cultural | 3.34 | 2.32 | 2.65 | 1.54 | 2.16 | 3.15 |
| Utility-centric | 1.10 | 0.89 | 0.93 | 0.65 | 1.38 | 1.30 |
| Recreational | 3.43 | 2.12 | 2.56 | 1.45 | 2.65 | 2.18 |
| Negative | 1.13 | 1.02 | 1.25 | 0.86 | 0.73 | 1.43 |

**Table 6.** Comparison of the three different scores for each of the nine attitudinal categories determined using MQDA, ChatGPT and Hugging Face transformers (using zero-shot classifier). Statistic used was Kruskal-Wallis rank sum test (df=2); * = *p* < 0.05, ** = *p* < 0.01, *** *p* < 0.005.

| **Attitudes** | ***T. khudree***<br>**($\chi^2$)** | ***T. putitora*(**<br>**$\chi^2$)** | ***T. remadevii***<br>**($\chi^2$)** | ***T. tambroide***<br>***s*($\chi^2$)** | ***T. tor***<br>**($\chi^2$)** | ***N. hexagono***<br>***lepis*($\chi^2$)** | **mahseer**<br>**($\chi^2$)** |
| --- | --- | --- | --- | --- | --- | --- | --- |
| Nature-centric | 0.83 | 1.72 | 1.46 | 0.45 | 0.16 | 1.94 | 2.04 |
| Ecological | 1.35 | 0.84 | 1.37 | 1.24 | 0.78 | 0.54 | 0.41 |
| Empathetic | 2.16 | 0.43 | 1.06 | 0.61 | 2.42 | 1.45 | 0.39 |
| Moralistic | 0.44 | 1.97 | 0.21 | 1.72 | 0.66 | 0.72 | 0.415 |
| Research-oriented | 0.64 | 2.76 | 0.34 | 0.54 | 0.13 | 0.84 | 0.37 |
| Religio-cultural | 1.32 | 0.11 | 2.04 | 1.23 | 0.87 | 1.07 | 1.47 |
| Utility-centric | 0.42 | 3.18 | 0.51 | 1.05 | 1.43 | 1.09 | 0.18 |
| Recreational | 0.42 | 1.54 | 0.24 | 0.91 | 0.85 | 1.04 | 0.73 |
| Negative | 1.65 | 2.05 | 1.14 | 2.15 | 2.40 | 1.87 | 2.16 |

**Table 7.** Inter-country and inter-species variation in each of the nine attitudinal categories (scores obtained using MQDA) derived from the mahseer news articles. Statistic used was Kruskal-Wallis rank sum test; * = *p* < 0.05, ** = *p* < 0.01, *** *p* < 0.005. NaN: not a number

| Attitudes | Inter-country | Inter-species |
| --- | --- | --- |
| | $\chi^2$<br>(df=5) | $\chi^2$<br>(df=6) |
| Nature-centric | 96.90*** | 177.93*** |
| Ecological | 33.77*** | 96.67*** |
| Empathetic | 2.28 | 306.56*** |
| Moralistic | 42.11*** | 205.24*** |
| Research-oriented | 24.32*** | 347.87*** |
| Religio-cultural | 37.61*** | 150.55*** |
| Utility-centric | 282.83*** | 172.60*** |
| Recreational | 47.05*** | 317.17*** |
| Negative | NaN | NaN |

**Fig 11.**
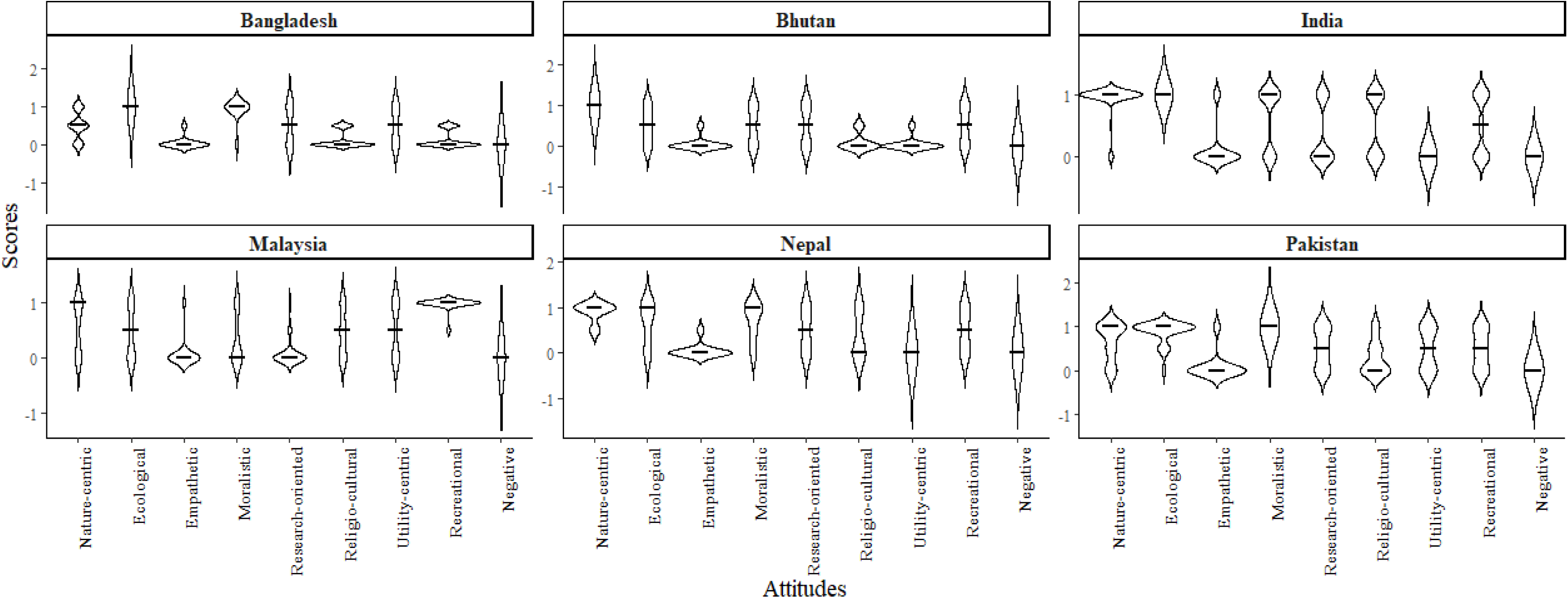
The attitudinal positioning scores of the news articles published from six focal countries. The horizontal line across the plot represents the median.

Positioning of the nine categories of attitudes in the mahseer related news were significantly different in all the six focal countries (Table 8a; Fig 11). Ecological and moralistic (over empathetic, recreational, religio-cultural, and negative; SM 11), and nature-centric and moralistic (empathetic, utility-centric and negative; SM 15) attitudes scored significantly high in Bangladeshi and Nepali newspapers, respectively. In India, ecological, nature-centric and empathetic attitudes were frequent with no significant differences found between the latter two (SM 13). Mahseer news from Pakistan also exhibited a significant dominance of moralistic and ecological attitudes (over empathetic, religio-cultural, utility-centric, recreational and negative SM 16). Here nature-centric were also significantly higher than both religio-cultural and negative but lesser than the empathetic. However, research-oriented, utility-centric and recreational attitudes scored significantly higher than empathetic attitudes. In Bhutan, nature-centric attitude was significantly higher than all other categories (SM 12). This nation‘s utility-centric attitude score was significantly greater than recreational but lesser than ecological and moralistic. Furthermore, the recreational, ecological and moralistic had a significantly higher representation than empathetic and negative, and the research-oriented scored higher only against the negative. Differing from other nations, recreational attitudes emerged as the most prominent, scoring significantly higher than all other categories in Malaysia, (SM 14). Interestingly, here the nature-centric and ecological attitudes scored significantly higher than empathetic, research-oriented and negative, and utility-centric and religio-cultural were significantly greater than both empathetic and research-oriented attitudes.

**Table 8.**
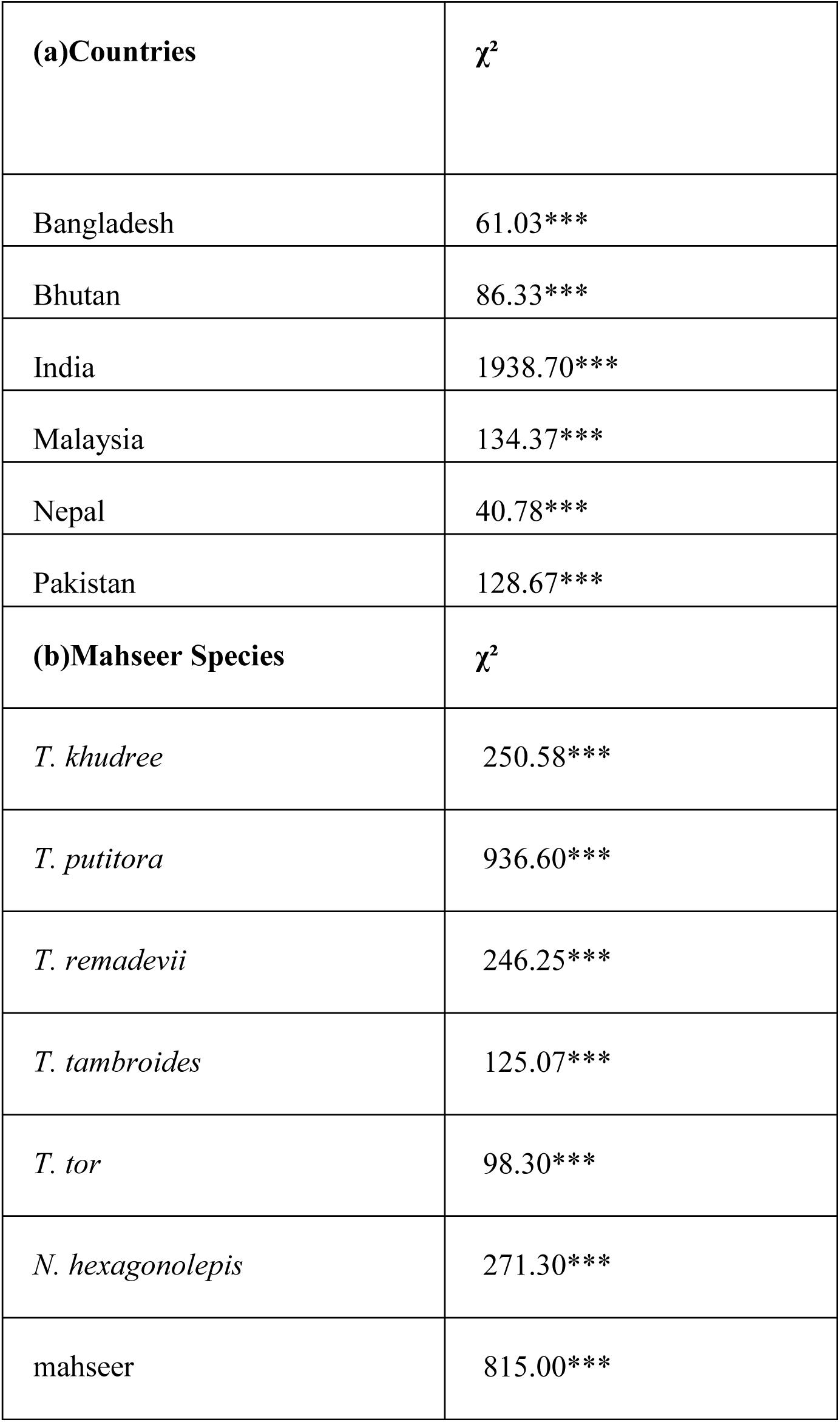
Inter-attitudinal category variation (scores obtained using MQDA) in the mahseer news from six focal countries (a) on different mahseer species (b). Statistic used was Kruskal-Wallis rank sum test (df=8); * = *p* < 0.05, ** = *p* < 0.01, *** *p* < 0.005.

Nature-centric attitude was significantly more prominent in *T. putitora* and ‗mahseer‘ (compared to *T. remadevii*, *T. tor, N. hexagonolepis* and *T. tambroides*; SM17) and *T. khudree* (than *T. remadevii*, *T. tambroides*, *T. tor* and *N. hexagonolepis*). Meanwhile, ecological dominated in the case of *T. remadevii*, *T. tor* and *N. hexagonolepis* (than all other species; SM 18) and moralistic in *T. khudree* and *T. remadevii* (compared to all other species SM 20). Research-oriented attitude prevaled news articles discussing *T. putitora*, *T. remadevii* and *T. tor* (than all other species SM 21) and so was the case of *T. khudree* and *N. hexagonolepis* when compared to *T. tambroides* and ‗mahseer‘. Religio-cultural attitude dominated in articles on *N. hexagonolepis* and ‗mahseer‘, (compared to all other species SM 22) and *T. tambroides* (than *T. khudree*, *T. putitora* and *T. remadevii*). The empathetic attitude, on the other hand, was most strongly expressed in news articles focusing on *T. tor* (than all other species SM 19). *T. khudree*, *T. remadevii*, *T. tambroides* and ‗mahseer‘ had more articles coming under this category than *T. putitora* and *N. hexagonolepis*. Both *N. hexagonolepis* and *T. tambroides* had significantly higher utility-centric score than *T. khudree*, *T. putitora*, *T. remadevii*, and ‗mahseer‘ SM 23). Similarly *T. putitora* and *T. tambroides* received significantly higher recreational attitude score than *T. khudree, T. remadevii*, *T. tor*, *N. hexagonolepis* and ‗mahseer‘ (SM 24). Interestingly, the negative attitude did not show any significant variation between species (Table 7; Fig 12).

**Fig 12.**
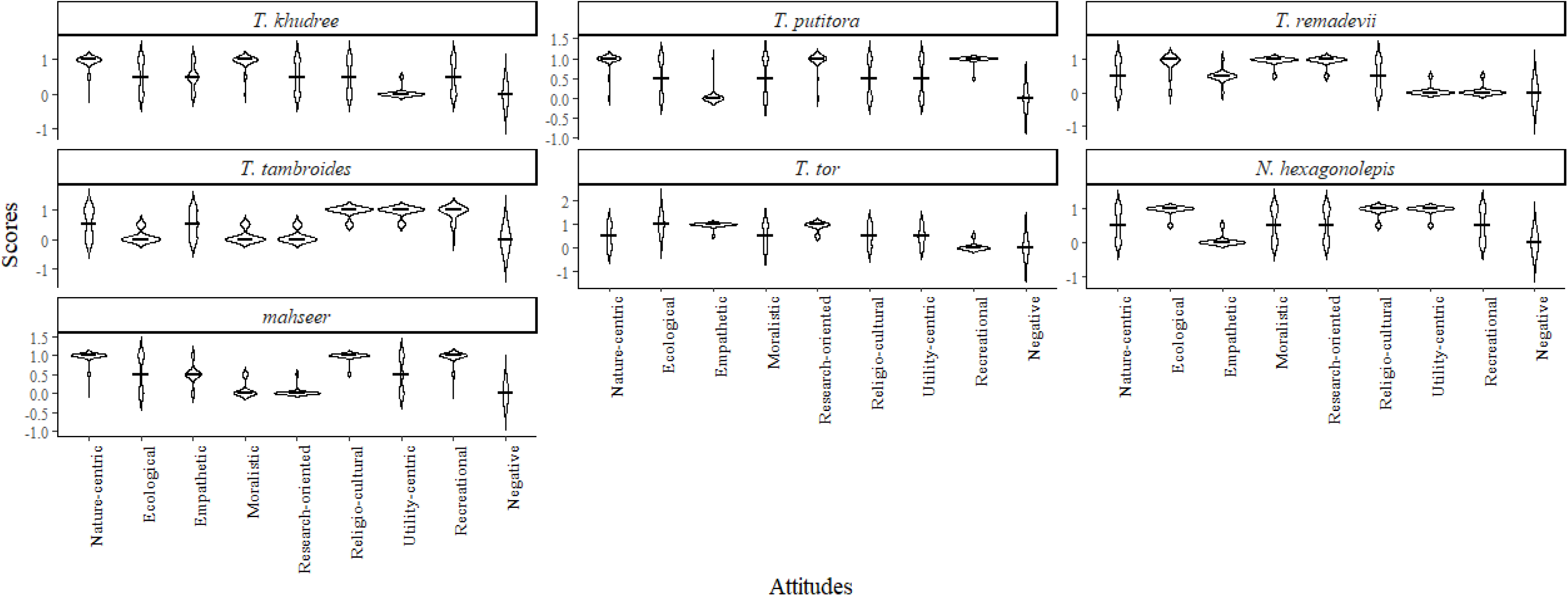
The attitudinal positioning scores of the articles reporting news on different mahseer species. The horizontal line across the plot represents the median

Attitudinal positioning scores received by the nine focal categories exhibited a significant intra-species difference (Table 8b; Fig 12). The nature-centric and moralistic, without any significant differences between, scored higher than all other categories in reports on *T. khudree* (SM 25). A greater presence of ecological and empathetic attitudes, in comparison to the utility-centric and negative, was also observed in the *T. khudree* related news. Higher nature-centric (compared to ecological, empathetic, moralistic, religio-cultural, utility-centric and negative; SM 26) and research-oriented (over ecological, moralistic, empathetic, utility-centric and negative) attitude scores were noticed in the case of *T. putitora.* A significant dominance of ecological and utility-centric (against nature-centric, empathetic, moralistic, research-oriented, recreational and negative ones SM 30) as well as the religio-cultural elements (higher than nature-centric, empathetic, moralistic, recreational and negative attitudes) made *N. hexagonolepis* distinct*. T. remadevii* received higher nature-centric (excluding empathetic and religio-cultural) and ecological (excluding moralistic and research-oriented) attitudes scores than all other categories (SM 27). *T. tor* news were mainly ecological (over moralistic, religio-cultural, utility-centric, recreational and negative ones SM 29) in nature. A different attitude positioning, with increased emphasis on recreational, religio-cultural and utility-centric attitudes (*T. tambroides* SM 28), was observed in the case of Malaysian mahseer species. Diverging from the species inhabiting South Asian nations‘ nature-centric attitude was only significantly higher than the negative in the case of these two species. A preference for nature-centric and religio-cultural attitudes (over all other categories except recreational) was observed in the news articles which did not specify the species of mahseer (category: ‗mahseer‘). Here, no significant difference was found between nature-centric and religio-cultural attitudes (SM 31).

## Discussion

Results of our study revealed that coverage of mahseer by the newspapers was higher in South Asian nations (India, Pakistan, Bangladesh, Bhutan and Nepal) than their South-East Asian counterparts. Of the 12 South-East Asian nations with reported mahseer populations news articles were available from only 3 - Malaysia, Indonesia and Thailand. However, the latter two published quite a low number (less than 5) of articles. A similar result obtained when the article collected were analysed keeping species in the focus; mahseers living in the rivers of South Asia (*T. putitora*, *T. khudree*, *T. tor*, *T. remadevii*, *N. hexagonolepis*), were represented more in the newspapers South-East Asian mahseer (*T. tambroides*). However, increase in the mahseer news articles published post 2015 from India may be associated with an array of activities conducted for conservation and commercial breeding during this period. A review of newspaper articles suggests that the observed increase is largely attributed to the establishment of a mahseer hatchery by the ICAR-Directorate of Coldwater Fisheries Research (DCFR) in Uttarakhand, India, in 2016 (ICAR-DCFR 2016). This was further amplified by the subsequent establishment of several other government hatcheries across various Indian states, including Himachal Pradesh (2016), Nagaland (2018), and Arunachal Pradesh (2019) (Gulati 2016; ICAR-DCFR 2018; Baruah et al. 2019). The newspaper articles also indicate that the hike in mahseer news observed during the time period 2021-2024 in India is linked to the Tata Power‘s ‗Act for Mahseer‘ project completing 50 years in 2021 (CSR Journal 2021). Within India, the states of Karnataka, Nagaland and the national capital New Delhi, emerged as major contributors of mahseer-related news reports. Karnataka is home to two of the well-known mahseer species, *T. khudree* and *T. remadevii*, while Nagaland harbours *T. putitora* and *N. hexagonolepis*, the latter being officially recognised as the State Fish. Furthermore, Karnataka is also home to two of the well-known recreational angling associations in India that have since decades, been associated with mahseer conservation works (Coorg Wildlife Society and Wildlife Association of South India) involving different stakeholder.

Among all species, *T. putitora* was the most prominently represented in the English newspapers, maybe due to its historical prominence as one of the most widely known, heavily angled and extensively studied mahseer species worldwide (Hamilton 1822; Bhatt and Pandit 2016; Pinder et al. 2019) Its broad geographical presence across countries like Pakistan, Nepal, Bhutan and Bangladesh, along with its presence in 10 Indian states and 2 Union Territories (UT), contributes to its media visibility. The species‘ designation as the National Fish of Pakistan and the State Fish of four Indian states (Himachal Pradesh, Uttarakhand, Arunachal Pradesh and UT of Jammu and Kashmir) further helps attract media attention and enhance its media appeal. Furthermore, launch of the ten year Golden Mahseer Conservation Action Plan (2022-2032) by the Bhutan in the year 2022 (Nature Conservation Division (NCD) 2022) and the presence of a national park dedicated to this species in Pakistan (Poonch River Mahseer National Park), further elevates the profile value of this fish among the conservation communicators. Attention of the policymakers, and the publication of the results of the scientific studies throwing light on important questions pertaining to the species also found to be improving media coverage on mahseers. For example, clarification of the taxonomic ambiguity of the humpback mahseer (*T. remadevii*) in 2018, along with its IUCN Red List classification (Bangalore Mirror Bureau 2018), and the revision of *T. mosal mahanadicus*, as a valid mahseer species, which was long considered to be a subspecies of *T. mosal* (Johnson et al. 2023), generated a number of newspaper articles in India. Similarly, declaration of different mahseer species as the state fish by different Indian states (*N. hexagonolepis* - Sikkim in 2021 (Sikkim Government Gazette 2021), *T. tor* - Madhya Pradesh in 2011 (MPSBB 2011) and *T. mahanadicus* - Odisha in 2012 (Mohanty et al. 2024)) was also seen reflecting as the increase in the number of news on mahseer. Nonetheless, the lower degree of coverage by the newspapers from the focal countries other than India and only *T. putitora*, getting their attention (that also to a certain extent only) exposes a major gap and raises concerns for the conservation communication strategies being followed for mahseer. Research has shown that news articles keep the potential to influence public discourse leading to the advocacy for policies or behaviour change supporting the protection of a species (King et al. 2017; Downing 2024). Therefore, it is vital to promote the interaction between mainstream print and visual media, and policymakers and researchers in all countries with wild populations of mahseer to ensure public support for the long-term plans to conserve this important group of freshwater fishes.

Newspapers from different nations framed headlines and content of mahseer news focusing mainly either environmental challenges and conservation, or the ecotourism and economy. In general, both headlines and text of the news from South Asian nations mostly projected conservation issues, environmental degradation, challenges faced, restocking efforts and cultural importance, while Malaysian media portrayed mahseer as food fish (culinary), sport fish (recreational angling) along with the associated ecotourism, travelling, commercial farming and revenue generation angles. This difference may be the result of the divergence present in public perception and priorities, cultural relationships and the importance given by academic community, media and policy makers to mahseers in their nations (Pinder et al. 2019; Sarma et al. 2022). For instance in Bhutan, *T. putitora* is considered to be one of the eight auspicious symbols in Buddhism, a religion widely practised in the nation (Philipp et al. 2015) and deep-rooted prominence given to the fishes in Bengali traditions (Roberts and Sen 1998) may have translating in to ‗cultural‘ frames commonly found in these area. Furthermore many nations in south Asia conduct active projects to conserve natural populations of different mahseers species in their freshwater ecosystems (Bhutan; Nature Conservation Division (NCD) 2022) and mahseer from this region *T. khudree*, *T. remadevii*, *T. tor* and *T. putitora* helping media to view them through the lens of ecological threat and conservation concern.

The prominence of economical angle in the Malaysian news and the mahseer from this region (*T. tambroides)* reveals how people from this nation consider mahseers not just as a species of ecological interest but also a contributor to their livelihood. This framing is aligned with the position of *T. tambroides* as the commercially valuable food fish in Malaysia. In this context it is also important to note that commercial breeding and the aquaculture of mahseer is significantly more prevalent in South-East Asia compared to the South Asian nations (Ramezani-Fard and Kamarudin 2012; Redhwan et al. 2022). Traditionally many South-East Asian nations (and a few communities of subcontinent) consider mahseers as a highly sought-after food fish and in high-end restaurants it fetches a very high rate. However, despite their popularity as culinary delicacies in Malaysia, the food-focused headlines were rare indicating a missed opportunity in promoting these species as culturally important cuisine and promoting in the tourism sector to support aquaculture. Furthermore, both headline and content frames of conservation, restocking, education and awareness and science and research were the lowest for *T. tambroides*. Although classified as data deficient (DD) by IUCN, the presence of natural populations of *T. tambroides* in different regions of Malaysia and the noticeable decline they are facing (Lau et al. 2021; Mulyani et al. 2022) demand public involved interventions to conserve them. Our results showing a higher aquaculture and economy-oriented and a lower ecological and research framing of the news on this species may lead to a higher fishing pressure on them in the wild. Framing of the headlines and matter of the news on another mahseer *N. hexagonolepis*, a species found in the selected regions of both South and Southeast Asia in terms of economic importance raises much concern. Such an action by the media may mask the urgent conservation needs of the near-threatened *N. hexagonolepis* whose wild populations are reportedly declining (Arunachalam 2010; Sarma et al. 2022). Hence, a more ecology centric and conservation focused journalism approach would help raise public awareness and create a pro-conservation narrative by highlighting these species‘ population decline and the need for conservation.

India stood out as a unique case in framing headlines and content of the mahseer news. Although the highest volume of the news on mahseer came from this nation, no single frame dominated either the headlines or the text. Such a diversity in the frames points towards a media or news fragmentation (increase in the number of available news media options with varied ideological leanings thereby creating an environment where users are confronted with an abundant choice of newspapers, each offering its own particular framing of news; Webster 2005; Steppat et al. 2023). Such a situation may lead to the dissolution of the ideas, splintering of the audience, possible dilution of the common and shared narrative and a segmented perception of reality (Knüpfer 2020), hindering the efforts done to converge public discourse around mahseers into a positive change in public attitude or policy to protect this fish. However, the presence of policy-related and human–human conflict angles in the headlines, though not available in the same ratio in the content, of the mahseer news from this country is a model for other nations. Although the content available in the text is also equally important, headlines mostly are the first element of a news article that gets registered in the mind of the readers. Studies have shown that headlines that evoke emotions, reduce the cognitive load and by being less informative increases curiosity and catches the reader‘s interest (Ifantidou 2023). They are the face of the news articles that draw the public into reading the story. This frame of the news also has an effect on what information the reader focuses on or ignores while going through the newspapers (McCrudden and Schraw 2007; Ecker et al. 2014). Interestingly, the headlines were predominantly neutral in the sentimental value they carried, across all focal nations and species. This result points towards a fact-based and objective reporting style with reduced elicitation of emotions in the case of mahseer. However, this style of communication promoting sentimental neutrality is a double edged sword; although it supports objective, unbiased, dissemination of the information such an approach could also reduce the emotional engagement necessary to stimulate public interest and policy demands in conservation narratives (Smith and Leiserowitz 2014; Myers et al. 2023). Surprisingly, in contrast to the content from other focal nations, Malaysian news articles and those focusing on *T. tambroides* were with more positive sentiment. Although, further analysis may be required to know the exact reason behind such a divergence, aquaculture, travel and tourism and other revenue generating activities centred on *T. tambroides* and other mahseer species popular in Malaysia may be persuading the journalists to choose more positive words during the communication.

Another feature of the mahseer news across all focal nations and species noted was the dominance of episodic framing - construction of the news based only on a specific event. Even though episodic frames dominate the news coverage worldwide across a multitude of topics (Tiegreen and Newman 2008), such a framing may lead readers to a simplistic view of a rather complex issue thereby preventing them from recognising the broader context and socio-cultural and ecological relevance of the information received (Iyengar 1991; Jacobson et al. 2012; Stafford et al. 2018; Correa-Chica et al. 2024). Hence, following such a news crafting strategy for data deficient species with declining population trends, a category which many mahseer species are currently positioned in is not an encouraging sign (Kottelat et al. 2018; Lau et al. 2021; Mulyani et al. 2022). Conversely framing the news in a thematic manner can offer more explanatory information and detailed knowledge about a topic, and positively influence the public‘s attitude and perception of that specific issue and make them aware of the role of various stakeholders in protecting mahseers and hold them responsible and accountable (Iyengar 1991; Siemer et al. 2007; Stafford et al. 2018; Correa-Chica et al. 2024). Hence, enhancing the awareness of journalists from the distribution range of mahseer about this bias existing in the framing of mahseer related news, and educating them to write the news giving proper context and population management shortcomings at the community and the system level rather than just mentioning a particular standalone incident is a need of the hour.

By including different stakeholders as messengers, the newspapers represent diverse perspectives popular in the society (Walker et al. 2019), as well leverage selected rhetoric and emotional appeal, establish credibility and authority (ethos, logos, pathos), and ultimately shape individual opinions, public discourse and narrative about mahseer conservation (Benoit-Barné et al. 2021; Correa-Chica et al. 2024; Gessa et al. 2024). Government officials were the most frequently quoted messengers in articles from India, Pakistan, Bangladesh, Bhutan and Malaysia and on all mahseer species. This result may be taken as the indicator of the government organisations leading the mahseer related activities in these geographical landscapes. Surprisingly, another nation from the same region Nepal showcased the much expected diversity in the voices from the society. Here the messengers included recreational anglers, celebrities such as authors, photographers and musicians along with military personnel revealing a broader cultural engagement with the masher. A similar trend found in the news articles from India and those focused on *T. putitora* and *T. remadevii* reiterate the potential for utilising the powerful voices of celebrities to achieve greater reach and influence regarding awareness (Omoyajowo et al. 2024) on mahseer amongst the public. *N. hexagonolepis* was unique; this fish received a relatively higher citation of politicians perhaps due to its image as a state fish in the states of Sikkim and Nagaland in India. Though less in number, the presence of temple and shrine caretakers as messengers in articles from India and Pakistan reflects the mahseer‘s religious and cultural significance this fish have in this region. Interestingly, news from most countries and about most species featuring local voices as messengers suggests an attempt to integrate grass-root level perspectives and the ground reality of the mahseer related activities happening in the society in their news by the media. This trend should be promoted since the words of the religious leaders (Myers et al. 2017), politicians (Agadjanian 2021) and peers from the community (Maki and Raimi 2017) may influence the public trust and acceptance of the news and direct the people‘s behaviours towards a positive direction for conservation (Whiting et al. 2019; Hollihan and Baaske 2022). However, the dearth of researchers in the role of messengers across all countries and mahseer species, except for news articles from Bangladesh and on *T. tor*, is of great concern. Although news from this nation and focusing on *T. tor* (species with higher proportions of education and awareness, and science and research frames) showed the least diversity in messengers, it had a higher number of mentions of researchers. Communicators working alongside scientists can help in eliminating misconceptions and misinformation prevalent in the communities related to the focal species, as well as promote a scientific approach towards ecological dilemmas involving those (Kusmanoff et al. 2020).

Along with selecting credible quoting sources, strategically positioning the news amongst the contents of the newspaper targeting diverse audiences and adding suitable visual representations are also important to achieve the objectives. Unfortunately, except in two South Asian nations (Nepal and Pakistan) and Malaysia (and the species from this nation, *T. tambroides*) English newspapers published very few mahseer related articles in the sections with the potential for persuasion (e.g. editorials, columns, letters, opinion pieces etc.; Benoit-Barné et al. 2021; Gessa et al. 2024). The relevance given by the society to mahseer can be seen reflecting as the number of persuasive articles (60%) published by the Nepali media. Since, information appearing in the sections of the newspapers such as editorials, columns, letters and opinion pieces carries a higher capability to influence the assimilation and accommodation of new information into the schema of a topic kept by the readers and their emotions (Benoit-Barné et al. 2021; Gessa et al. 2024; White 2025), communicators from nations other than Nepal, and especially Bangladesh with a count of zero article belonging to this category, needs to be made aware of the importance of including mahseer news in the influential columns.

It is well-known that ―pictures speak a thousand stories‖ and research has proven that two specific visual frames - solution/action and threat/problem oriented – can influence the pro-conservation behaviours (O‘Neill and Nicholson-Cole 2009; Metag et al. 2016; Ison et al. 2024). A conservation action-oriented frame can evoke the sense of hope, make conservation issues more tangible to the public and announce that solutions are achievable, and thereby stimulate pro-conservation intentions (Metag et al. 2016; Ison et al. 2024. Meanwhile, the problem/threat-oriented ones evoke feelings of helplessness by giving a connotation of ‗the issue is too big to solve‘. Such visuals can distance people from the real issues and may lead to the complete disengagement with the text available (Chadwick 2015; Ison et al. 2024). Unfortunately, except Indian and Malaysian media none of the focal nations gave the emphasis it requires to the visual representation in their articles on mahseer and a total absence of the image was a surprising feature of the mahseer news from Bangladesh. However, the absence of any dominant visual frame, despite this high representation, suggests a lack of effective visual framing strategy for mahseers in Indian newspapers and for *T. putitora* the most visually represented species. Furthermore, amongst the images extracted those related to the conservation action (research, monitoring, breeding, awareness and education) were the least in comparison to the conservation threat/problem frames. A manual exploration of the images collected from the news articles revealed that the conservation action related ones mainly came from the South Asian nations and focused on their native mahseer species. Furthermore, a large share of images used by the communicators was the aesthetically appealing portrayal of the species and landscapes (rivers and forests etc.). While such visually pleasing images may attract reader‘s attention, they often fail to represent and convey the depth of the challenges faced by the focal species (Ison et al. 2024). The underrepresentation of the conservation action-oriented and stakeholder-related images across focal nations and species highlights a significant gap and a possibly missed opportunity for more effective public engagement. Currently, most of the newspapers maintain their online presence, and with the popularisation of digital technologies, internet and social media availability of quality images are also not rare. Editors of the newspapers from the focal nations should utilise these opportunities to increase the impact of their mahseer news.

The attitude positioning echoed the pattern of framing observed in the headlines and the content of the news, when inter and intra-nation as well as species specific comparisons were conducted. Newspapers from South Asia generously promoted nature-centric and ecological attitudes reflecting their positive affinity for nature, environmental and ecological concerns and conservation issues. The news articles coming from the category ‗mahseer‘ also projecting nature-centric and religio-cultural attitudes reveals that when mahseer is discussed without specific taxonomic context, discourse tends to elaborate on the cultural and environmental significance of these fishes. Furthermore, higher level of religio-cultural and moralistic attitude positioning observed in the articles from India and Pakistan respectively, reiterate the cultural value these fishes have in these region (Gupta et al. 2016; Pinder et al. 2019) and point towards the moral and ethical obligations towards nature and biodiversity prevalent in the conservation narratives of this region (Gupta and Guha 2002; Shivaramakrishnan 2015). The empathetic attitudes received relatively a lower score for several countries other than India, and *T. tor* was most strongly associated with this kind of attitude, although this species is a target of recreational anglers across the globe (Das and Binoy 2024). The dominance of recreational and utility-centric attitudes reaffirms the unique socio-economic and leisure based media positioning of mahseer in Malaysia with respect to angling, tourism, farming, food and revenue and other values attributed to this fish beyond conservation. In this nation tourists are allowed hold mahseer, feel their bites feed them and play with without hurting and enjoy mahseer foot and body massage at the Mahseer Sanctuaries or red-zones established by the Tagal Community Management Systems (community-based fisheries resource management) under the Fisheries Department (DOF), (Wong et al. 2009; Lee et al. 2022). In this context it is also important to note that Malaysian news articles included relatively more fish farmers, restaurant and resort owners as messengers. Hence, mahseer aquaculture and ecotourism popular in Malaysia may be the catalyst behind the type of attitude positioning observed here.

Along with studying the framing and attitude positioning in the news related to mahseers, the present study also explored the potential of Artificial Intelligence (AI) based tools in analysing the large amount of text generated by multiple media and integrating diverse information to get insights on conservation communication. In this era of information explosion data management is a major issue faced by communication researchers and GenAI based tools are often advised as a solution (Şen et al. 2023). Our results showing negligible variation in the results of framing and attitudinal positioning analysis conducted using popular Large Language Model (ChatGPT) and Natural Language Processing tools (zero-shot Hugging Face transformers), in comparison to the conventionally used manual coding (MQDA) for these purposes support the growing trend of human-AI collaboration in thematic QDA (Bijker et al. 2024; Perkins and Roe 2024; Prescott et al. 2024; Yan et al. 2024). Extending these lines of analysis to the conservation communication focusing on freshwater ecosystems and the biodiversity it harbours can expedite the evidence based planning and policy making for protecting these ecosystems providing multiple vital services. Although such GenAI algorithms can significantly reduce the time and resources required for conducting analysis of large amounts of text, care should be taken at each and every step of data cleaning, uploading and interpreting the result. In many contexts these general purpose tools may function as ‗black boxes‘ and the errors arising at any step of the data processing can lead to wrong interpretations with unexpected consequences (Perkins and Roe 2024). So performance of AI tools should be validated with the type of data and the context of the study in mind and the development of frames and their final interpretation should be based on the researchers‘ in-depth, contextual, critical and analytical viewpoints (Bijker et al. 2024; Christou 2024).

Our study is not devoid of limitations. Lack of availability of a large number of news articles from the focal nations other than India and the use of only common and scientific names of the mahseer species for surfing might have impacted the data availability. In nations where English-language newspapers are not popular, selecting this language may target only a selected section of the population and may even be aimed at composing news with the international community in mind (Holland 2006). Such newspapers may not give importance or space to the problems having impact only at the local level. Including the vernacular newspapers like Hindi, Bangla, Urdu, Nepalese, Malay, Thai, etc. would have provided a more elucidated picture of mahseer communication happening at the grassroot level.

## Conclusion

The newspapers and journalists are key actors in making people aware of the conservation status, research conducted, cultural and traditional importance of any species, including mahseers, along with a myriad of ecosystem services provided by them. The information shared by newspapers also influences the perceptions, sets attitudes and direct actions of the public and gradually shapes social norms for conserving a species or an ecosystem. The present study revealed that newspapers from South Asia and the articles focusing on the mahseer species native to this region followed framing that project conservation issue, environmental degradation, challenges faced, restocking efforts and cultural importance of these fishes in their headlines, text and images chosen. The English newspapers from this region attempted attitude positioning to promote engagement in pro-conservation behaviours. By contrast, Malaysian newspapers portrayed mahseer through a utilitarian lens emphasising their role as food and sport fish and highlighted commercial farming, ecotourism, travel and revenue generation perspectives. However, although this divergence seen between the nations located at the South and South Eastern regions of Asia may be the reflection of cultural, social and economic importance given to mahseer by the local communities, a communication approach highlighting not only these fishes‘ ecological and conservation importance, but also elaborating its socio-economic values and the ecosystem services provided by them, might evoke more obligation from the public to protect them (Kusmanoff et al. 2020). Such an integrated approach rightly blending these two popular perspectives is a need of the hour since natural populations of mahseer present in the Malaysia and neighbouring nations requires conservation support (Lau et al. 2021; Mulyani et al. 2022) and many South Asian nations considers mahseers as a delicacy and conducts programmes to popularise their aquaculture (Barbhuiya et al. 2013; Jha et al. 2019). However, interventions to bring such changes in communication strategies need to be done with caution as any sudden and drastic shift in the popular frames used by the media may lead to the ignorance of the information by the public (Berk 2025).

The news coverage, framing strategy and attitude positioning is also influenced by the ideology followed by the publishers and editorial culture present in the newspaper (Painter 2013; O‘Neill 2020). Hence, along with studying different qualities of the materials focusing on mahseer published by the newspapers (including those in the vernacular languages) from different nations and understanding their political and ideological views is also essential to get the ground realities of mahseer conservation communication. In this context it is also important to note that, disseminating under-contextualised and oversimplified information can dilute the efforts done through communication to conserve a species or an ecosystem (Jacobson et al. 2012; Muter et al. 2013; Chandelier et al. 2018; López-Baucells et al. 2023). Our study revealed that newspapers from many focal nations seldom framed mahseer news projecting scientific and educational elements, promoted an attitude supporting it, and rarely included researches as the messengers. In order to promote trustworthy and inclusive conservation communication along with strengthening interaction and collaboration between scientists and journalists (Bagla and Binoy 2017), enhancing the incorporation of the voices of diverse representatives from the society such as locals, NGOs, government officials are vital (Kusmanoff 2017; Kusmanoff et al. 2020). Our result also highlights the possibilities of using GenAI and NLP based tools as collaborative agents in conservation communication research. This approach will significantly reduce the time and resources required for analysing big data to provide vital insights for the ecosystem managers and policy makers, in comparison to the conventional manual analysis. Hence future analysis of different forms of traditional media, social media and other data sources popularly utilised by the iEcologists and conservation culturomics researchers, integrated with the exploration of the determinants of the public acceptance of conservation communication may help in establishing effective information dissemination strategies to bring positive behavioural change at the individual level and translating it into injunctive social norms (behaviours that one is expected to follow and expects others to follow; Farrow et al. 2017; Kusmanoff et al. 2020) and public-driven policies to protect the natural populations of the ‗Tiger of the Rivers‘ and their habitats worldwide.

## Author Contributions

**Prantik Das:** Conceptualisation; methodology; investigation; data curation; formal analysis; visualisation; writing - original draft preparation; writing - review and editing. **V. V. Binoy:** Conceptualisation; methodology; visualisation; writing - review and editing; supervision.

## Supporting information

Supplementary Materials. Das and Binoy 2025

## Acknowledgements

Prantik acknowledges the Human Research Development Group (HRDG) - Council of Scientific and Industrial Research (CSIR), New Delhi (09/1320(0001)/2020-EMR-I) for the research fellowship.

## Data Availability Statement

The data will be made available upon request from the corresponding author.

