## Supplementary Materials. Das and Binoy 2025 for "‘Crafting Fishy News’: Framing and Attitudinal Positioning in English Newspaper Articles on Mahseer from Their Endemic Range"

**Supplementary Materials (SM)**

SM 1. Scientific and common names of the focal mahseer species used as the keywords to search the newspapers

| **S. No.** | **Scientific Name** | **Common Name**  (Vattakaven et al. 2016: India Biodiversity Portal; Froese and Pauly 2025: FishBase) |
| --- | --- | --- |
| 1 | *Tor putitora* | Golden Mahseer; Himalayan Mahseer |
| 2 | *T. khudree* | Deccan Mahseer; Blue-finned Mahseer |
| 3 | *T. remadevii* | Humpback Mahseer; Orange-finned Mahseer |
| 4 | *T. mosal* | Mosal Mahseer |
| 5 | *T. tor* | Tor Mahseer; Deep-bodied Mahseer; Red-finned  Mahseer |
| 6 | *T. barakae* | Barak Mahseer |
| 7 | *T. mahanadicus* | Mahanadi Mahseer |
| 8 | *T. malabaricus* | Malabar Mahseer |
| 9 | *T. kulkarnii* | Dwarf Mahseer |
| 10 | *T. mussullah* | Mussullah Mahseer |
| 11 | *T. ater* | Dark Mahseer |
| 12 | *T. dongnaiensis* | Dongnai Mahseer |
| 13 | *T. mekongensis* | Mekong Mahseer |
| 14 | *T. tambra* | Javan Mahseer; Red Mahseer |
| 15 | *T. laterivittatus* | Dark-striped Mahseer |
| 16 | *T. polylepis* | — |
| 17 | *T. sinensis* | Chinese Mahseer; Red Mahseer |
| 18 | *T. tambroides* | Thai Mahseer; Malay Mahseer; Malaysian Mahseer |
| 19 | *T. douronensis* | Pink Mahseer; Pink Brook Carp |
| 20 | *T. yingjiangensis* | Yingjiang Mahseer |
| 21 | *T. macrolepis* | Indus Mahseer |
| 22 | *T. progenius* | Jungha Mahseer |
| 23 | *Neolissochilus hexagonolepis* | Chocolate Mahseer; Copper Mahseer |
| 24 | *N. hexastichus* | — |
| 25 | *N. wynaadensis* | Wayanad Mahseer |
| 26 | *N. soro* | — |
| 27 | *N. hemispinus* | — |
| 28 | *N. dukai* | — |
| 29 | *N. spinulosus* | Spinulosus Mahseer |
| 30 | *N. acutirostris* | — |
| 31 | *N. capudelphinus* | — |
| 32 | *N. micropthalmus* | — |
| 33 | *N. minimus* | — |
| 34 | *N. pnar* | Cave Mahseer; Blind Mahseer |
| 35 | *N. tamiraparaniensis* | — |
| 36 | *N. kaladanensis* | — |
| 37 | *N. stracheyi* | — |
| 38 | *N. blanci* | — |
| 39 | *N. paucisqaumatus* | — |
| 40 | *N. soroides* | Soro Brook Carp or Antimony Fish |
| 41 | *N. subterraneus* | Cave Brook Carp |
| 42 | *N. longipinnis* | — |
| 43 | *N. nigrovittatus* | — |
| 44 | *N. thienemanni* | — |
| 45 | *N. benasi* | — |
| 46 | *N. stevensonii* | Stevenson’s Masheer |
| 47 | *N. vittatus* | — |
| 48 | *N. baoshanensis* | — |
| 49 | *N. blythii* | — |
| 50 | *N. compressus* | — |
| 51 | *N. hendersoni* | — |
| 52 | *N. sumatranus* | Sumatran Mahseer |
| 53 | *N. innominatus* | — |
| 54 | *N. qiaojiensis* | — |
| 55 | *Naziritor chelynoides* | Dark Mahseer |
| 56 | *Naziritor zhobensis* | Zhobi or Zhob Mahseer |

SM 2.  List of nations and mahseer species with fewer than 10 newspaper articles (these were not considered for analyses).

| **Countries** | **Number of Newspaper Articles** |
| --- | --- |
| China | 2 |
| Indonesia | 3 |
| Thailand | 4 |
| **Species** | **Number of Newspaper Articles** |
| *T. douronensis* | 1 |
| *T. sinensis* | 1 |
| *T. tambra* | 7 |
| *N. soroides* | 4 |
| *N. wynaadensis* | 2 |

SM 3. *Post hoc* analysis of nature-centric attitude in mahseer news across all six focal nations. Statistics used was Dunn test with Bonferroni method. *** =  p<0.001, ** =  p<0.01, * = p<0.05

|  | **Bhutan**  **(Z)** | **India**  **(Z)** | **Malaysia**  **(Z)** | **Nepal**  **(Z)** | **Pakistan**  **(Z)** |
| --- | --- | --- | --- | --- | --- |
| **Bangladesh** | -5.56*** | -6.09 *** | -2.00 | -3.54 | -2.53 |
| **Bhutan** | - | 1.08 | 4.88  *** | 1.33 | 4.08  *** |
| **India** | - | - | 6.42  *** | 0.84 | 4.96  *** |
| **Malaysia** | - | - | - | -2.37 | -0.78 |
| **Nepal** | - | - | - | - | 1.80 |

SM 4. *Post hoc* analysis of ecological attitude in mahseer news across all six focal nations. Statistics used was Dunn test with Bonferroni method. *** =  p<0.001, ** =  p<0.01, * = p<0.05

|  | **Bhutan**  **(Z)** | **India**  **(Z)** | **Malaysia**  **(Z)** | **Nepal**  **(Z)** | **Pakistan**  **(Z)** |
| --- | --- | --- | --- | --- | --- |
| **Bangladesh** | 2.22 | 2.13 | 4.43  *** | 2.19 | 1.11 |
| **Bhutan** | - | -0.82 | 2.37 | 0.32 | -1.51 |
| **India** | - | - | 4.94  *** | 0.98 | -1.32 |
| **Malaysia** | - | - | - | -1.50 | -4.49  *** |
| **Nepal** | - | - | - | - | -1.54 |

SM 5. *Post hoc* analysis of empathetic attitude in mahseer news across all six focal nations. Statistics used was Dunn test with Bonferroni method. *** =  p<0.001, ** =  p<0.01, * = p<0.05

|  | **Bhutan**  **(Z)** | **India**  **(Z)** | **Malaysia**  **(Z)** | **Nepal**  **(Z)** | **Pakistan**  **(Z)** |
| --- | --- | --- | --- | --- | --- |
| **Bangladesh** | -0.13 | -0.55 | 0.19 | -0.11 | -0.21 |
| **Bhutan** | - | -0.50 | 0.39 | 0.00 | -0.09 |
| **India** | - | - | 1.30 | 0.36 | 0.48 |
| **Malaysia** | - | - | - | -0.30 | -0.55 |
| **Nepal** | - | - | - | - | -0.07 |

SM 6. *Post hoc* analysis of moralistic attitude in mahseer news across all six focal nations. Statistics used was Dunn test with Bonferroni method. *** =  p<0.001, ** =  p<0.01, * = p<0.05

|  | **Bhutan**  **(Z)** | **India**  **(Z)** | **Malaysia**  **(Z)** | **Nepal**  **(Z)** | **Pakistan**  **(Z)** |
| --- | --- | --- | --- | --- | --- |
| **Bangladesh** | 2.63 | 2.18 | 3.80  ** | 1.19 | -0.52 |
| **Bhutan** | - | -1.41 | 1.09 | -1.16 | -3.97  ** |
| **India** | - | - | 3.65  ** | -0.40 | -4.38  *** |
| **Malaysia** | - | - | - | -2.11 | -5.91  *** |
| **Nepal** | - | - | - | - | -1.89 |

SM 7. *Post hoc* analysis of research-oriented attitude in mahseer news across all six focal nations. Statistics used was Dunn test with Bonferroni method. *** =  p<0.001, ** =  p<0.01, * = p<0.05

|  | **Bhutan**  **(Z)** | **India**  **(Z)** | **Malaysia**  **(Z)** | **Nepal**  **(Z)** | **Pakistan**  **(Z)** |
| --- | --- | --- | --- | --- | --- |
| **Bangladesh** | 0.39 | 1.61 | 3.19* | 0.24 | 0.15 |
| **Bhutan** | - | 1.44 | 3.31* | -0.10 | -0.31 |
| **India** | - | - | 3.44  ** | -1.15 | -2.25 |
| **Malaysia** | - | - | - | -2.68 | -4.14  *** |
| **Nepal** | - | - | - | - | -0.14 |

SM 8. *Post hoc* analysis of religio-cultural attitude in mahseer news across all six focal . Statistics used was Dunn test with Bonferroni method. *** =  p<0.001, ** =  p<0.01, * = p<0.05

|  | **Bhutan**  **(Z)** | **India**  **(Z)** | **Malaysia**  **(Z)** | **Nepal**  **(Z)** | **Pakistan**  **(Z)** |
| --- | --- | --- | --- | --- | --- |
| **Bangladesh** | 0.00 | -3.61  ** | -2.79 | -1.90 | -1.79 |
| **Bhutan** | - | -4.62  *** | -3.34 | -2.10 | -2.12 |
| **India** | - | - | 0.75 | 0.76 | 2.41 |
| **Malaysia** | - | - | - | 0.32 | 1.31 |
| **Nepal** | - | - | - | - | 0.57 |

SM 9. *Post hoc* analysis of utility-centric attitude in mahseer news across all six focal nations. Statistics used was Dunn test with Bonferroni method. *** =  p<0.001, ** =  p<0.01, * = p<0.05

|  | **Bhutan**  **(Z)** | **India**  **(Z)** | **Malaysia**  **(Z)** | **Nepal**  **(Z)** | **Pakistan**  **(Z)** |
| --- | --- | --- | --- | --- | --- |
| **Bangladesh** | 5.75  *** | 7.17  *** | 0.40 | 4.90  *** | 0.02 |
| **Bhutan** | - | 0.00 | -7.09  *** | 0.00 | -7.29  *** |
| **India** | - | - | -11.49  *** | 0.00 | -11.25  *** |
| **Malaysia** | - | - | - | 5.52  *** | -0.51 |
| **Nepal** | - | - | - | - | -5.75  *** |

SM 10. *Post hoc* analysis of recreational attitude in mahseer news across all six focal nations. Statistics used was Dunn test with Bonferroni method. *** =  p<0.001, ** =  p<0.01, * = p<0.05

|  | **Bhutan**  **(Z)** | **India**  **(Z)** | **Malaysia**  **(Z)** | **Nepal**  **(Z)** | **Pakistan**  **(Z)** |
| --- | --- | --- | --- | --- | --- |
| **Bangladesh** | -2.44 | -3.07 * | -5.78  *** | -2.09 | -2.62 |
| **Bhutan** | - | -0.03 | -3.71  *** | -0.01 | 0.00 |
| **India** | - | - | -5.97  *** | 0.01 | 0.04 |
| **Malaysia** | - | - | - | 2.88 | 4.23  *** |
| **Nepal** | - | - | - | - | 0.01 |

SM 11. *Post hoc* analysis of the nine different categories of attitudes present in the mahseer news from Bangladesh. Statistics used was Dunn test with Bonferroni method. *** = p<0.001, ** =  p<01, * = p<0.05

|  | **Ecological**  **(Z)** | **Empathetic**  **(Z)** | **Moralistic**  **(Z)** | **Research-oriented**  **(Z)** | **Religio-cultural**  **(Z)** | **Utility-centric**  **(Z)** | **Recreational**  **(Z)** | **Negative**  **(Z)** |
| --- | --- | --- | --- | --- | --- | --- | --- | --- |
| **Nature-centric** | -2.45 | 2.61 | -2.01 | 0.21 | 2.10 | 0.14 | 2.10 | 2.86 |
| **Ecological** | — | 5.06*** | 0.44 | 2.66 | 4.55*** | 2.59 | 4.55*** | 5.32*** |
| **Empathetic** | — | — | -4.62*** | -2.40 | -0.51 | -2.47 | -0.51 | 0.26 |
| **Moralistic** | — | — | — | 2.22 | 4.11*** | 2.15 | 4.11*** | 4.88*** |
| **Research-oriented** | — | — | — | — | -1.89 | -0.07 | -1.89 | -2.66 |
| **Religio-cultural** | — | — | — | — | — | -1.96 | 0.00 | -0.77 |
| **Utility-centric** | — | — | — | — | — | — | -1.96 | -2.73 |
| **Recreational** | — | — | — | — | — | — | — | -0.77 |

SM 12. *Post hoc* analysis of the nine different categories of attitudes present in the mahseer news from Bhutan. Statistics used was Dunn test with Bonferroni method. *** =  p<0.001, ** =  p<0.01, * = p<0.05

|  | **Ecological**  **(Z)** | **Empathetic**  **(Z)** | **Moralistic**  **(Z)** | **Research-oriented**  **(Z)** | **Religio-cultural**  **(Z)** | **Utility-centric**  **(Z)** | **Recreational**  **(Z)** | **Negative**  **(Z)** |
| --- | --- | --- | --- | --- | --- | --- | --- | --- |
| **Nature-centric** | 3.24* | 6.57*** | 3.32* | 3.41* | 6.13*** | 6.57*** | 3.32* | 7.01*** |
| **Ecological** | — | 3.33* | 0.09 | 0.18 | 2.89 | 3.33* | 0.09 | 3.77* |
| **Empathetic** | — | — | -3.24* | -3.15 | -0.44 | 0.00 | -3.24* | 0.44 |
| **Moralistic** | — | — | — | 0.09 | 2.80 | 3.24* | 0.00 | 3.68* |
| **Research-oriented** | — | — | — | — | -2.71 | 3.15 | 0.09 | 3.59* |
| **Religio-cultural** | — | — | — | — | — | 0.44 | 2.80 | 0.88 |
| **Utility-centric** | — | — | — | — | — | — | 3.24* | 0.44 |
| **Recreational** | — | — | — | — | — | — | — | 3.68* |

SM 13. *Post hoc* analysis of the nine different categories of attitudes present in the mahseer news from India. Statistics used was Dunn test with Bonferroni method. *** =  p<0.001, ** =  p<0.01, * = p<0.05

|  | **Ecological**  **(Z)** | **Empathetic**  **(Z)** | **Moralistic**  **(Z)** | **Research-oriented**  **(Z)** | **Religio-cultural**  **(Z)** | **Utility-centric**  **(Z)** | **Recreational**  **(Z)** | **Negative**  **(Z)** |
| --- | --- | --- | --- | --- | --- | --- | --- | --- |
| **Nature-cnetric** | -2.22 | 23.92*** | 9.02*** | 17.06*** | 12.16*** | 27.65*** | 12.53*** | 27.65*** |
| **Ecological** | - | 26.13*** | 11.24*** | 19.28*** | 14.38*** | 29.87*** | 14.75*** | 29.87*** |
| **Empathetic** | - | - | -14.90*** | -6.86*** | -11.75*** | 3.73*** | -11.39*** | 3.73*** |
| **Moralistic** | - | - | - | 8.04*** | 3.14 | 18.63*** | 3.51*** | 18.63 *** |
| **Research-oriented** | - | - | - | - | 4.90*** | 10.59*** | 4.53*** | 10.59*** |
| **Religio-cultural** | - | - | - | - | - | 15.49*** | 0.37 | 15.49*** |
| **Utility-centric** | - | - | - | - | - | - | -15.12*** | 0.00 |
| **Recreational** | - | - | - | - | - | - | - | 15.12*** |

SM 14. *Post hoc* analysis of the nine different categories of attitudes present in the mahseer news from Malaysia. Statistics used was Dunn test with Bonferroni method. *** =  p<0.001, ** =  p<0.01, * = p<0.05

|  | **Ecological**  **(Z)** | **Empathetic**  **(Z)** | **Moralistic**  **(Z)** | **Research-oriented**  **(Z)** | **Religio-cultural**  **(Z)** | **Utility-centric**  **(Z)** | **Recreational**  **(Z)** | **Negative**  **(Z)** |
| --- | --- | --- | --- | --- | --- | --- | --- | --- |
| **Nature-centric** | 1.24 | 4.99*** | 2.24 | 4.79*** | 1.08 | 0.80 | -3.53* | 5.49*** |
| **Ecological** | - | 3.78*** | 1.00 | 3.57* | -0.17 | -0.45 | -4.81*** | 4.28*** |
| **Empathetic** | - | - | -2.77 | -0.21 | -3.94*** | -4.22*** | -8.59*** | 0.50 |
| **Moralistic** | - | - | - | 2.56 | -1.17 | -1.45 | -5.81*** | 3.28* |
| **Research-oriented** | - | - | - | - | 3.73 1*** | -4.01*** | -8.38*** | 0.71 |
| **Religio-cultural** | - | - | - | - | - | -0.28 | -4.64*** | 4.45*** |
| **Utility-centric** | - | - | - | - | - | - | -4.36*** | 4.72*** |
| **Recreational** | - | - | - | - | - | - | - | 9.09*** |

SM 15. *Post hoc* analysis of the nine different categories of attitudes present in the mahseer news from Nepal. Statistics used was Dunn test with Bonferroni method. *** = p<0.001, ** = p<0.01, * = p<0.05

|  | **Ecological**  **(Z)** | **Empathetic**  **(Z)** | **Moralistic**  **(Z)** | **Research-oriented**  **(Z)** | **Religio-cultural**  **(Z)** | **Utility-centric**  **(Z)** | **Recreational**  **(Z)** | **Negative**  **(Z)** |
| --- | --- | --- | --- | --- | --- | --- | --- | --- |
| **Nature-centric** | 1.23 | 4.10*** | 0.96 | 1.91 | 2.47 | 4.38*** | 1.91 | 4.38*** |
| **Ecological** | - | 2.87 | -0.28 | 0.68 | 1.23 | 3.14 | 0.68 | 3.14 |
| **Empathetic** | - | - | -3.14* | -2.19 p=1.00 | -1.63 p=1.00 | 0.28 p=1.00 | -2.19 p=1.00 | 0.28 p=1.00 |
| **Moralistic** | - | - | - | 0.96 | 1.51 | 3.42* | 0.96 | 3.42* |
| **Research-oriented** | - | - | - | - | -0.55 | 2.47 | 0.00 | -2.47 |
| **Religio-cultural** | - | - | - | - | - | 1.91 | 0.55 | -1.91 |
| **Utility-centric** | - | - | - | - | - | - | 2.47 | 0.00 |
| **Recreational** | - | - | - | - | - | - | - | -2.47 |

SM 16. *Post hoc* analysis of the nine different categories of attitudes present in the mahseer news from Pakistan. Statistics used was Dunn test with Bonferroni method. *** =  p<0.001, ** =  p<0.01, * = p<0.05

|  | **Ecological**  **(Z)** | **Empathetic**  **(Z)** | **Moralistic**  **(Z)** | **Research-oriented**  **(Z)** | **Religio-cultural**  **(Z)** | **Utility-centric**  **(Z)** | **Recreational**  **(Z)** | **Negative**  **(Z)** |
| --- | --- | --- | --- | --- | --- | --- | --- | --- |
| **Nature-centric** | 1.78 | -4.95*** | 2.74 | 1.37 | 3.29* | 1.51 | 1.51 | 5.77*** |
| **Ecological** | - | 6.73*** | -0.96 | 3.15 | 5.08*** | 3.29* | 3.29* | 7.55*** |
| **Empathetic** | - | - | -7.69*** | -3.58* | -1.65 | -3.44* | -3.44* | 0.82 |
| **Moralistic** | - | - | - | 4.11*** | 6.03*** | 4.25*** | 4.25*** | 8.51*** |
| **Research-oriented** | - | - | - | - | -1.92 | 0.14 | -0.14 | 4.40*** |
| **Religio-cultural** | - | - | - | - | - | -1.78 | 1.79 | 2.48 |
| **Utility-centric** | - | - | - | - | - | - | 0.00 | 4.26*** |
| **Recreational** | - | - | - | - | - | - | - | 4.26*** |

SM 17. *Post hoc* analysis of nature-centric attitude in mahseer news across all mahseer species. Statistics used was Dunn test with Bonferroni method. *** =  p<0.001, ** =  p<0.01, * = p<0.05

|  | ***T. putitora***  **(Z)** | ***T. remadevii***  **(Z)** | ***T. tambroides***  **(Z)** | ***T. tor***  **(Z)** | ***N. hexagonolepis***  **(Z)** | **mahseer**  **(Z)** |
| --- | --- | --- | --- | --- | --- | --- |
| ***T. khudree*** | -0.63 | 6.29  *** | 4.61  ** | 4.58  *** | 6.55  *** | 0.72 |
| ***T. putitora*** | - | 7.93  *** | 5.49  *** | 5.40  ** | 8.67  *** | 0.18 |
| ***T. remadevii*** | - | - | -0.42 | -0.27 | 0.36 | -7.70  *** |
| ***T. tambroides*** | - | - | - | 0.12 | 0.15 | -5.43  *** |
| ***T. tor*** | - | - | - | - | 0.01 | -5.35  *** |
| ***N. hexagonolepis*** | - | - | - | - | - | -8.31  *** |

SM 18. *Post hoc* analysis of ecological attitude in mahseer news across all mahseer species. Statistics used was Dunn test with Bonferroni method. *** =  p<0.001, ** =  p<0.01, * = p<0.05

|  | ***T. putitora***  **(Z)** | ***T. remadevii***  **(Z)** | ***T. tambroides***  **(Z)** | ***T. tor***  **(Z)** | ***N. hexagonolepis***  **(Z)** | **mahseer**  **(Z)** |
| --- | --- | --- | --- | --- | --- | --- |
| ***T. khudree*** | -0.30 | -3.68* | 3.85  ** | -3.96  ** | -5.04  *** | 0.04 |
| ***T. putitora*** | - | -4.07  ** | 4.44  ** | -4.16  *** | -5.91  *** | 0.34 |
| ***T. remadevii*** | - | - | 6.38  *** | -1.05 | 0.87 | 4.11  ** |
| ***T. tambroides*** | - | - | - | -6.35  *** | -7.43  *** | -4.17  ** |
| ***T. tor*** | - | - | - | - | 0.43 | 4.21  *** |
| ***N. hexagonolepis*** | - | - | - | - | - | 5.81  *** |

SM 19. *Post hoc* analysis of empathetic attitude in mahseer news across all mahseer species. Statistics used was Dunn test with Bonferroni method. *** =  p<0.001, ** =  p<0.01, * = p<0.05

|  | ***T. putitora***  **(Z)** | ***T. remadevii***  **(Z)** | ***T. tambroides***  **(Z)** | ***T. tor***  **(Z)** | ***N. hexagonolepis***  **(Z)** | **mahseer**  **(Z)** |
| --- | --- | --- | --- | --- | --- | --- |
| ***T. khudree*** | 9.23  *** | -0.50 | 0.37 | -3.91  ** | 7.42  *** | 0.32 |
| ***T. putitora*** | - | -8.32  *** | -5.42  *** | -9.84  *** | -0.20 | -12.53  *** |
| ***T. remadevii*** | - | - | 0.72 | -3.32* | 7.09  *** | 0.28 |
| ***T. tambroides*** | - | - | - | -3.53  * | 4.97  *** | 0.61 |
| ***T. tor*** | - | - | - | - | 9.01  *** | 3.98  ** |
| ***N. hexagonolepis*** | - | - | - | - | - | -8.93  *** |

SM 20. *Post hoc* analysis of moralistic attitude in mahseer news across all mahseer species. Statistics used was Dunn test with Bonferroni method. *** =  p<0.001, ** =  p<0.01, * = p<0.05

|  | ***T. putitora***  **(Z)** | ***T. remadevii***  **(Z)** | ***T. tambroides***  **(Z)** | ***T. tor***  **(Z)** | ***N. hexagonolepis***  **(Z)** | **mahseer**  **(Z)** |
| --- | --- | --- | --- | --- | --- | --- |
| ***T. khudree*** | 5.80  *** | -0.37 | 7.02  *** | 3.25* | 4.76  *** | 11.51  *** |
| ***T. putitora*** | - | -5.29  *** | 4.09  *** | 0.06 | 0.24 | 8.45  *** |
| ***T. remadevii*** | - | - | 6.85  *** | 3.32* | 4.60  *** | 10.23  *** |
| ***T. tambroides*** | - | - | - | -2.91 | 3.52* | 0.02 |
| ***T. tor*** | - | - | - | - | 0.08 | 3.74  ** |
| ***N. hexagonolepis*** | - | - | - | - | - | 5.66  *** |

SM 21. *Post hoc* analysis of research-oriented attitude in mahseer news across all mahseer species. Statistics used was Dunn test with Bonferroni method. *** =  p<0.001, ** =  p<0.01, * = p<0.05

|  | ***T. putitora***  **(Z)** | ***T. remadevii***  **(Z)** | ***T. tambroides***  **(Z)** | ***T. tor***  **(Z)** | ***N. hexagonolepis***  **(Z)** | **mahseer**  **(Z)** |
| --- | --- | --- | --- | --- | --- | --- |
| ***T. khudree*** | -6.20  *** | -4.76  *** | 3.17 * | -3.42* | 0.23 | 6.40  *** |
| ***T. putitora*** | - | -0.41 | 7.41  *** | -0.01 | 5.72  *** | 17.03  *** |
| ***T. remadevii*** | - | - | 6.55  *** | 0.25 | 4.48  *** | 10.82  *** |
| ***T. tambroides*** | - | - | - | -5.36  *** | -3.30* | 0.78 |
| ***T. tor*** | - | - | - | - | 3.22* | 7.67  *** |
| ***N. hexagonolepis*** | - | - | - | - | - | 6.48  *** |

SM 22. *Post hoc* analysis of religio-cultural attitude in mahseer news across all mahseer species. Statistics used was Dunn test with Bonferroni method. *** =  p<0.001, ** =  p<0.01, * = p<0.05

|  | ***T. putitora***  **(Z)** | ***T. remadevii***  **(Z)** | ***T. tambroides***  **(Z)** | ***T. tor***  **(Z)** | ***N. hexagonolepis***  **(Z)** | **mahseer**  **(Z)** |
| --- | --- | --- | --- | --- | --- | --- |
| ***T. khudree*** | -0.01 | -0.03 | -3.95  ** | -0.16 | -5.60  *** | -7.21  *** |
| ***T. putitora*** | - | -0.02 | -4.36  *** | -0.17 | -6.87  *** | -10.02  *** |
| ***T. remadevii*** | - | - | -3.68  ** | -0.14 | -4.95  *** | -6.11  *** |
| ***T. tambroides*** | - | - | - | 3.02 | 0.11 | 0.46 |
| ***T. tor*** | - | - | - | - | -3.71  ** | -4.34  *** |
| ***N. hexagonolepis*** | - | - | - | - | - | 0.51 |

SM 23. *Post hoc* analysis of utility-centric attitude in mahseer news across all mahseer species. Statistics used was Dunn test with Bonferroni method. *** =  p<0.001, ** =  p<0.01, * = p<0.05

|  | ***T. putitora***  **(Z)** | ***T. remadevii***  **(Z)** | ***T. tambroides***  **(Z)** | ***T. tor***  **(Z)** | ***N. hexagonolepis***  **(Z)** | **mahseer**  **(Z)** |
| --- | --- | --- | --- | --- | --- | --- |
| ***T. khudree*** | -6.91  *** | -0.06 | -7.65  *** | -3.91  ** | -10.84  *** | -6.64  *** |
| ***T. putitora*** | - | 5.72  *** | -4.09  ** | -0.11 | -6.65  *** | 0.21 |
| ***T. remadevii*** | - | - | -7.12  *** | -3.63  ** | -9.58  *** | -5.60  *** |
| ***T. tambroides*** | - | - | - | 2.87 | 0.22 | 3.88  ** |
| ***T. tor*** | - | - | - | - | -3.64  ** | -0.02 |
| ***N. hexagonolepis*** | - | - | - | - | - | 6.12  *** |

SM 24. *Post hoc* analysis of recreational attitude in mahseer news across all mahseer species. Statistics used was Dunn test with Bonferroni method. *** =  p<0.001, ** =  p<0.01, * = p<0.05

|  | ***T. putitora***  **(Z)** | ***T. remadevii***  **(Z)** | ***T. tambroides***  **(Z)** | ***T. tor***  **(Z)** | ***N. hexagonolepis***  **(Z)** | **mahseer**  **(Z)** |
| --- | --- | --- | --- | --- | --- | --- |
| ***T. khudree*** | -8.51  *** | 5.05  *** | -3.75  ** | 3.84  ** | -0.06 | -7.36  *** |
| ***T. putitora*** | - | 13.07  *** | 1.23 | 9.33  *** | 8.31  *** | 0.88 |
| ***T. remadevii*** | - | - | -7.31  *** | -0.07 | -4.91  *** | -11.96  *** |
| ***T. tambroides*** | - | - | - | 6.17  *** | 3.75  ** | -0.79 |
| ***T. tor*** | - | - | - | - | -3.75  ** | -8.72  *** |
| ***N. hexagonolepis*** | - | - | - | - | - | -7.23  *** |

SM 25. *Post hoc* analysis of the nine different categories of attitudes present in *T. khudree* news. Statistics used was Dunn test with Bonferroni method. *** =  p<0.001, ** =  p<0.01, * = p<0.05

|  | **Ecological**  **(Z)** | **Empathetic**  **(Z)** | **Moralistic**  **(Z)** | **Research-oriented**  **(Z)** | **Religio-cultural**  **(Z)** | **Utility-centric**  **(Z)** | **Recreational**  **(Z)** | **Negative**  **(Z)** |
| --- | --- | --- | --- | --- | --- | --- | --- | --- |
| **Nature-centric** | 4.73*** | 5.18*** | 0.00 | 5.22*** | 5.32*** | 10.96*** | 5.32*** | 11.36*** |
| **Ecological** | - | 0.45 | -4.73*** | 0.49 | 0.59 | 6.23*** | 0.59 | 6.63*** |
| **Empathetic** | - | - | -5.18*** | 0.04 | 0.14 | 5.78*** | 0.14 | 6.18*** |
| **Moralistic** | - | - | - | 5.22*** | 5.32*** | 10.96*** | 5.32*** | 11.36*** |
| **Research-oriented** | - | - | - | - | -0.10 | 5.74*** | -0.10 | 6.14*** |
| **Religio-cultural** | - | - | - | - | - | 5.64*** | -0.00 | 6.04*** |
| **Utility-centric** | - | - | - | - | - | - | 5.64*** | -0.40 |
| **Recreational** | - | - | - | - | - | - | - | 6.04*** |

SM 26. *Post hoc* analysis of the nine different categories of attitudes present in *T. putitora* news. Statistics used was Dunn test with Bonferroni method. *** =  p<0.001, ** =  p<0.01, * = p<0.05

|  | **Ecological**  **(Z)** | **Empathetic**  **(Z)** | **Moralistic**  **(Z)** | **Research-oriented**  **(Z)** | **Religio-cultural**  **(Z)** | **Utility-centric**  **(Z)** | **Recreational**  **(Z)** | **Negative**  **(Z)** |
| --- | --- | --- | --- | --- | --- | --- | --- | --- |
| **Nature-centric** | 8.51*** | 18.98*** | 8.91*** | 0.77 | 10.15*** | 9.77*** | -0.59 | 20.30*** |
| **Ecological** | - | 10.47*** | 0.40 | -7.74*** | 1.64 | 1.26 | -9.11*** | 11.79*** |
| **Empathetic** | - | - | -10.07*** | -18.21*** | -8.83  *** | -9.21  *** | -19.57  *** | 1.33 |
| **Moralistic** | - | - | - | -8.14  *** | 1.24 | 0.86 | -9.51 *** | 11.39  *** |
| **Research-oriented** | - | - | - | - | -0.38  *** | 9.00  *** | 1.36 | 19.53  *** |
| **Religio-cultural** | - | - | - | - | - | -0.38 | 10.74  *** | 10.15  *** |
| **Utility-centric** | - | - | - | - | - | - | 10.37  *** | 10.53  *** |
| **Recreational** | - | - | - | - | - | - | - | 20.90  *** |

SM 27. *Post hoc* analysis of the nine different categories of attitudes present in *T. remadevii* news. Statistics used was Dunn test with Bonferroni method. *** =  p<0.001, ** =  p<0.01, * = p<0.05

|  | **Ecological**  **(Z)** | **Empathetic**  **(Z)** | **Moralistic**  **(Z)** | **Research-oriented**  **(Z)** | **Religio-cultural**  **(Z)** | **Utility-centric**  **(Z)** | **Recreational**  **(Z)** | **Negative**  **(Z)** |
| --- | --- | --- | --- | --- | --- | --- | --- | --- |
| **Nature-centric** | 3.73  *** | 0.08 | 4.60  *** | -4.47  *** | 0.00 | 4.64  *** | 4.64 *** | 5.02  *** |
| **Ecological** | - | 3.67  *** | -0.87 | -0.75 | 3.73  *** | 8.37  *** | 8.37  *** | 8.75  *** |
| **Empathetic** | - | - | -4.55  *** | -4.43  *** | 0.08 | 4.75  *** | 4.75  *** | 5.13  *** |
| **Moralistic** | - | - | - | 0.12 | 4.60  *** | 9.24  *** | 9.24  *** | 9.62  *** |
| **Research-oriented** | - | - | - | - | 4.47  *** | 9.12  *** | 9.12  *** | 9.50  *** |
| **Religio-cultural** | - | - | - | - | - | 4.64  *** | 4.64  *** | 5.02  *** |
| **Utility-centric** | - | - | - | - | - | - | 0.00 | 0.38 |
| **Recreational** | - | - | - | - | - | - | - | 0.38 |

SM 28. *Post hoc* analysis of the nine different categories of attitudes present in *T. tambroides* news. Statistics used was Dunn test with Bonferroni method. *** =  p<0.001, ** =  p<0.01, * = p<0.05

|  | **Ecological**  **(Z)** | **Empathetic**  **(Z)** | **Moralistic**  **(Z)** | **Research-oriented**  **(Z)** | **Religio-cultural**  **(Z)** | **Utility-centric**  **(Z)** | **Recreational**  **(Z)** | **Negative**  **(Z)** |
| --- | --- | --- | --- | --- | --- | --- | --- | --- |
| **Nature-centric** | 3.05 | 0.13 | 3.05 | 3.05 | -2.87 | -2.87 | -2.53 | 3.79  *** |
| **Ecological** | - | -2.92 | 0.00 | 0.00 | -5.92 *** | -5.92  *** | -5.58  *** | 0.74 |
| **Empathetic** | - | - | 2.92 | 2.92 | -2.99 | -2.99 | -2.65 | 3.66  *** |
| **Moralistic** | - | - | - | 0.00 | -5.92  *** | -5.92  *** | -5.58 *** | 0.74 |
| **Research-Oriented** | - | - | - | - | 5.92  *** | -5.92  *** | 5.58 *** | 0.74 |
| **Religio-cultural** | - | - | - | - | - | 0.00 | -0.34 | 6.65  *** |
| **Utility-centric** | - | - | - | - | - | - | 0.34 | 6.65  *** |
| **Recreational** | - | - | - | - | - | - | - | 6.31  *** |

SM 29. *Post hoc* analysis of the nine different categories of attitudes present in *T. tor* news. Statistics used was Dunn test with Bonferroni method. *** =  p<0.001, ** =  p<0.01, * = p<0.05

|  | **Ecological**  **(Z)** | **Empathetic**  **(Z)** | **Moralistic**  **(Z)** | **Research-oriented**  **(Z)** | **Religio-cultural**  **(Z)** | **Utility-centric**  **(Z)** | **Recreational**  **(Z)** | **Negative**  **(Z)** |
| --- | --- | --- | --- | --- | --- | --- | --- | --- |
| **Nature-centric** | 3.43* | 3.23* | 0.16 | -2.84 | -0.13 | 0.10 | 3.00 | 3.33* |
| **Ecological** | - | 0.19 | 3.27* | 0.58 | 3.30* | 3.53* | 6.43  *** | 6.75  *** |
| **Empathetic** | - | - | 3.07 | 0.39 | 3.11 | 3.33* | 6.24  *** | 6.56  *** |
| **Moralistic** | - | - | - | -2.68 | 0.03 | 0.26 | 3.16 | 3.49* |
| **Research-oriented** | - | - | - | - | -2.72 | 2.94 | 5.85  *** | 6.17  *** |
| **Religio-cultural** | - | - | - | - | - | 0.23 | 3.13 | 3.45* |
| **Utility-centric** | - | - | - | - | - | - | 2.90 | 3.22* |
| **Recreational** | - | - | - | - | - | - | - | 0.32 |

SM 30. *Post hoc* analysis of the nine different categories of attitudes present in *N. hexagonolepis* news. Statistics used was Dunn test with Bonferroni method. *** =  p<0.001, ** =  p<0.01, * = p<0.05

|  | **Ecological**  **(Z)** | **Empathetic**  **(Z)** | **Moralistic**  **(Z)** | **Research-oriented**  **(Z)** | **Religio-cultural**  **(Z)** | **Utility-centric**  **(Z)** | **Recreational**  **(Z)** | **Negative**  **(Z)** |
| --- | --- | --- | --- | --- | --- | --- | --- | --- |
| **Nature-centric** | -4.98  *** | 5.42  *** | 0.04 | 0.06 | -4.76  *** | -4.98  *** | 0.46 | 5.88  *** |
| **Ecological** | - | 10.41  *** | 5.02*** | 5.04  *** | 0.22 | 0.00 | 5.44  *** | 10.86  *** |
| **Empathetic** | - | - | -5.39  *** | -5.37  *** | -10.19  *** | -10.41  *** | -4.97  *** | 0.46 |
| **Moralistic** | - | - | - | 0.02 | -4.80  *** | -5.02  *** | 0.42 | 5.84  *** |
| **Research-oriented** | - | - | - | - | 4.82  *** | -5.04  *** | -0.40 | -5.83  *** |
| **Religio-cultural** | - | - | - | - | - | -0.22 0 | 5.22  *** | 10.65  *** |
| **Utility-centric** | - | - | - | - | - | - | 5.44  *** | 10.86  *** |
| **Recreational** | - | - | - | - | - | - | - | 5.42  *** |

SM 31. *Post hoc* analysis of the nine different categories of attitudes present in ‘mahseer’ category news. Statistics used was Dunn test with Bonferroni method. *** =  p<0.001, ** =  p<0.01, * = p<0.05

|  | **Ecological**  **(Z)** | **Empathetic**  **(Z)** | **Moralistic**  **(Z)** | **Research-oriented**  **(Z)** | **Religio-cultural**  **(Z)** | **Utility-centric**  **(Z)** | **Recreational**  **(Z)** | **Negative**  **(Z)** |
| --- | --- | --- | --- | --- | --- | --- | --- | --- |
| **Nature-centric** | 7.64  *** | 8.41  *** | 15.54  *** | 16.81  *** | -0.07 | 8.24  *** | 0.34 | 17.27  *** |
| **Ecological** | - | 0.77 | 7.90  *** | 9.17  *** | -7.70  *** | 0.61 | -7.30  *** | 9.63  *** |
| **Empathetic** | - | - | 7.13  *** | 8.40  *** | -8.47  *** | -0.16 | -8.07  *** | 8.86  *** |
| **Moralistic** | - | - | - | 1.26 | -15.61  *** | -7.30  *** | -15.21  *** | 1.73 |
| **Research-oriented** | - | - | - | - | -16.87  *** | 8.56  *** | -16.47  *** | 0.47 |
| **Religio-cultural** | - | - | - | - | - | 8.31  *** | -0.40 | 17.34  ** |
| **Utility-centric** | - | - | - | - | - | - | -7.91  *** | 9.03  *** |
| **Recreational** | - | - | - | - | - | - | - | 16.93  *** |

____________________________
